# Catestatin improves heart metabolic flexibility by promoting mitochondrial structure and function

**DOI:** 10.1101/2024.12.29.630689

**Authors:** Venkat R. Chirasani, Satadeepa Kal, Nilima Biswas, Sumana Mahata, Kechun Tang, Teresa Pasqua, Ennio Avolio, Suborno Jati, Gautam Bandyopadhyay, Brian P. Head, Hemal H. Patel, Geert van den Bogaart, Debashis Sahoo, Sanjib Senapati, Sushil K. Mahata

## Abstract

Hypertension, a major cause of cardiomyopathy, is one of the most critical risk factors for heart failure and mortality worldwide. Loss of metabolic flexibility of cardiomyocytes is one of the major causes of heart failure. Although Catestatin (CST) treatment is known to be both hypotensive and cardioprotective, its effect on cardiac metabolism is unknown. In this study, we undertook a transcriptomic approach to identify differentially expressed genes that were filtered using Boolean implication relationships to develop a model of gene regulation in saline or CST-supplemented CST knockout (CST-KO) mice. The analysis revealed a set of gene signatures (fibroblast, cardiomyocyte, and macrophage) rescued after CST supplemented CST-KO mice compared to wild-type. Furthermore, we independently validated these gene signature models using publicly available patient datasets. Since the gene signature includes genes related to glucose, fatty acid metabolism, and mitochondrial function, we assessed the glucose and fatty acid uptake after CST treatment. We found that CST treatment can restore the cardiac metabolic inflexibility in CST-KO heart due to the metabolic shift of glucose utilization to fatty acid as energy source. Binding studies after immunoprecipitation and mass spectrometry revealed CST binding with ATP synthase, supported by molecular simulation and computational modeling that predicted CST binding to α/β subunit of ATP synthase. Colocalization of CST with mitochondria and increased mitochondrial membrane potential and ATP production upon CST treatment in neonatal cardiomyocytes further exhibit CST as a key regulator of cardiac metabolism and mitochondrial function.

## Introduction

The heart has the highest oxygen demand per tissue mass (4.3 mmol/kg*min) and is uniquely adapted to generate 6 to 25 kg of adenosine triphosphate (ATP) daily to fuel its contractile apparatus and ionic pumps ^1-3^. In the normal adult heart, mitochondrial oxidative phosphorylation contributes to almost all (>95%) of ATP production. Since the heart stores relatively low levels of ATP (0.5 µmol/g wet weight), it requires continuous and rapid production of new ATP molecules since the myocardial ATP pool is consumed in 10 s ^4^. Therefore, to sustain sufficient ATP production, the heart acts as an “omnivore” ^5^ and utilizing multiple energy-producing substrates to produce ATP: triglycerides and non-esterified (free) fatty acids (FFAs; contributing to 60% to 90% ATP FFA β-oxidation), carbohydrates (glucose and lactate; contributing to 10% to 30% ATP production) and to some extent also ketone bodies and amino acids (contributing to 5% to 10% ATP production) ^1^. The normal heart has the innate capacity to switch between different energy substrates, referred to as “metabolic flexibility” ^6^. Thus, during injury/stress, the heart shifts from using FFAs as energetic substrates toward glucose. This metabolic flexibility is compromised with the development of insulin resistance, called “metabolic inflexibility,” when the β-oxidation of FFAs becomes the main source of myocardial energy production^6^.

We have previously shown that Chromogranin A (CgA)-derived peptide Catestatin (CST: hCgA_352-372_) ^7^ acts as an anti-hypertensive ^8-10^ and improves insulin sensitivity in diet-induced obese (DIO) mice by reducing inflammation ^11^, decreasing endoplasmic reticulum stress and inhibiting gluconeogenesis ^11, 12^. We have recently found CST knockout (CST-KO) mice, lacking the 60 bp of the *CHGA* gene coding for CST, are resistant to insulin on a normal chow diet ^11^. Additionally, CST is cardioprotective, because CST-KO are not only hypertensive but also display left ventricular hypertrophy and have elevated levels of proinflammatory cytokines and catecholamines, which can be reversed by administration of exogenous CST ^13^.

Insulin receptors are abundantly expressed in cardiomyocytes, where insulin promotes glucose and fatty acid uptake, but inhibits the use of FFAs as an energy source ^14-16^. Insulin resistance is associated with increased plasma levels of FFAs, which leads to increased uptake of FFAs and reduction in insulin-stimulated glucose uptake, resulting in shifting of ATP production in the heart almost exclusively from beta-oxidation of FFAs ^14-16^. It has been shown that the altered substrate preference precedes the development of cardiac dysfunction, implicating altered cardiac metabolism in the development of diabetic cardiomyopathy ^17, 18^. Therefore, we hypothesized that the cardioprotective effects of CST are caused by improved metabolic flexibility of the cardiomyocytes.

To test this hypothesis, we used machine learning to identify differentially expressed genes (DEGs) that were filtered using Boolean implication relationships ^19, 20^ to develop a model of gene regulation in the absence (CST-KO) and presence of CST (WT and CST-KO supplemented with CST). Since we found increased fibrosis and macrophage infiltration in CST-KO heart ^13^, we focused the Boolean analysis on genes specifically expressed in fibroblasts, cardiomyocytes and macrophages. In addition, we made independent validation of CST gene signature model using publicly available human (GSE141910, GSE5406, GSE194297, GSE107480) and mice (GSE241771) datasets. We found that CST directly located to the mitochondria, where it could bind to ATP-synthase, promoting ATP synthesis and the mitochondrial membrane potential. Thus, CST is a key regulator of heart metabolic flexibility.

## Material and Methods

### Animals

Male WT and CST-KO (20-24 weeks old) on C57BL/6 were studied. Animals were kept in a 12 hr dark/light cycle and fed a normal chow diet (NCD: 14% calorie from fat; LabDiet 5P00). All studies on animals were approved by the UCSD and Veteran Affairs San Diego Institutional Animal Care and Use Committees (IACUC) and conform to relevant National Institutes of Health guidelines.

### RNA Sequencing

Total RNA was isolated from left ventricle of C57BL6 WT (4), CST-KO (4) and CST-KO+CST mice (4) using RNeasy kit (Qiagen). RNA was quantified using Nanodrop spectrophotometer (Thermo Scientific^TM^), and integrity was evaluated using Tapestation (Agilent). The following complementary DNA library preparation and sequencing were performed at the UCSD IGM core facility.

### Boolean analysis of RNA Seq data to assess CST regulation of gene expression in the heart

We employed a composite score, as previously described ^20^, to construct a model of gene regulation in the absence of CST (CST-KO) and in the presence of CST (WT and CST-KO supplemented with CST). Given the observed increase in fibrosis and macrophage infiltration in CST-KO hearts, we focused on genes specifically expressed in fibroblasts, cardiomyocytes, and macrophages, utilizing Boolean analysis ^19^.

For this analysis, we leveraged a large dataset comprising 25,955 human samples (GSE119087), profiled using Affymetrix Human U133 Plus 2.0 microarrays. To identify fibroblast- and cardiomyocyte-specific genes, we applied the Boolean Implication relationship DCN low => X low. Boolean Implication relationships are discovered by using two parameters Sthr and pThr as described before ^19^. Sthr = 3.0 and pthr=0.1 is used to discover these Boolean Implication relationships which has a false discovery rate < 1e-4.

Using these gene sets, we developed an algorithm to select a subset of genes for a composite score that adhered to the desired gene regulatory relationships in the absence (CST-KO) and presence of CST (WT and CST-KO supplemented with CST). The CST gene signature models were independently validated using publicly available human (GSE141910, GSE5406, GSE194297, GSE107480) and mouse (GSE241771) datasets.

### *In vivo* tissue glucose uptake and metabolism

*In vivo* glucose uptake and production of glucose-6-phosphate (G6P) and glycogen was assessed following a published protocol^21^ but using double isotopes -^3^H-glucose and ^14^C-2-deoxyglucose (2DG) as described previously by us ^22^.

### *In vivo* fatty acid incorporation and lipid extraction and oxidation

A 100 µl solution of U-^14^C-palmitic acid-BSA complex (molar ratio 2.5:1), containing 1 µCi and 250 µM palmitate was injected intraperitoneally per mouse. After 90 min, mice were sacrificed, blood was saved, and tissues were subjected to lipid extraction by Bligh and Dyer’s adaptation ^23^ of Folch’s original method ^24^. Radioactivity in the lower chloroform layer was determined to measure fatty acids and glycogen.

### Protein analysis by immunoblotting

Left ventricles were homogenized in RIPA lysis buffer containing phosphatase and protease inhibitors, as previously described ^25^. Homogenates were subjected to SDS-PAGE and immunoblotted. Primary antibodies for phosphorylated and total AKT, and GSK-3β were from Cell Signaling Technology.

### Real Time PCR

Total RNA from the heart was isolated by using the RNeasy Mini Kit and reverse-transcribed using the qScript cDNA synthesis kit. cDNA samples were amplified using PERFECTA SYBR FASTMIX L-ROX 1250 and analyzed on an Applied Biosystems 7500 Fast Real-Time PCR system. All PCRs were normalized to *Rplp0*, and relative expression levels were determined by the ΔΔ*C_t_* method.

### Transmission Electron Microscopy (TEM)

To displace blood and wash tissues before fixation, mice were cannulated through the apex of the heart and perfused with a Ca^2+^ and Mg^2+^ free buffer composed of DPBS, and 9.46 mM KCl (to arrest hearts in diastole), as described previously ^22^. Fixation, embedding, sectioning and staining were done following our previous publication ^22^. Grids were viewed using a JEOL 1200EX II TEM and photographed using a Gatan digital camera.

### Morphometric analysis

Sarcomere length in micrographs was determined using ImageJ software. Cristae membrane surface area was also measured using ImageJ by manually tracing each crista within a mitochondrion. The cristae surface was then normalized by dividing the sum of the total cristae membrane surface area with the outer membrane surface area per mitochondrion, as described previously ^22^.

### CST modeling

CST is an endogenous cationic peptide, comprising of 21 amino acids and is hydrophobic in nature. CST exhibits potent catecholamine release-inhibitory activity by acting on the neuronal nicotinic acetylcholine receptor ^7, 26^. The structure of human wild-type CST was determined by NMR (PDB ID: 1LV4) ^27^. However, the absence of any secondary structural elements has prompted the researchers to find a high-resolution conformation of this peptide ^28^. In this work, we have generated a 3D structure of CST following a similar protocol as proposed earlier ^28^. The NMR structure of CST was downloaded from protein data bank and subjected to short energy minimization for optimal positioning of the side chains. The minimized structure was subsequently subjected to 200 nanoseconds explicit water molecular dynamics (MD) simulations to generate an ensemble of CST conformations.

### F1-ATP synthase modeling

ATP synthase is one of the most abundant enzymes in living organisms. It plays a crucial role in synthesizing ATP, the energy currency of cells. ATP synthase is composed of two constituent subunits F0 and F1. Our LC-MS/MS and TOF data suggest that CST specifically binds to the α-β interface of F1-ATP synthase. Hence, we modeled mouse F1-ATP synthase based on the available crystal structure of bovine F1-ATP synthase (PDB ID:1E79) ^29^. A 97% similarity between two sequences provided us a very accurate model of mouse F1-ATP synthase, obtained through homology modelling. MODELLER9v13 ^30^ was used to generate the initial model structure. Two ATP and three ADP were included in the structure, as similar to the bovine F1-ATP synthase crystal conformation. An extensive set of minimization, thermalization, and finally a long 100 ns MD simulation was performed on this structure to obtain a more sophisticated model of F1-ATP synthase for protein-protein docking.

### Protein-protein docking of CST on F1-ATP synthase

Protein-protein docking *of CST on F1-ATP synthase* was performed using HEX docking algorithm ^31^ to locate the probable binding site of CST in F1-ATP synthase structure. During molecular docking, CST was allowed to explore the entire F1-ATP synthase structure and identify the best binding region (unbiased docking). The best docked complex was identified based on HEX docking score. This structure of CST-F1-ATP synthase complex was also refined by molecular dynamics (MD) simulation for 100 ns in explicit water.

### MD Simulation methodology

All the simulations were performed using Gromacs-4.5.5 simulation package ^32^ and CHARMM36 force-field parameters ^33, 34^. First, the system was energy minimized using conjugate gradient and steepest descent algorithms, each with 1000 steps. The energy-minimized structure was subsequently solvated in a cubic periodic box with about 180,000 explicit water molecules. TIP3P model was used to describe the water molecules. The salt concentration of 0.15 M was maintained with appropriate number of Na^+^ and Cl^-^ ions to mimic the physiological ionic strength. The solvated system was again energy minimized and subsequently heated to 310 K. The system was equilibrated in NPT ensemble at 1 atm and 310 K for about 10ns, by when the solvent density, system temperature reached to a plateau. This followed a production phase of 100 ns, on which all analyses were performed. Long-range electrostatic interactions were described using Particle Mesh Ewald sum technique with a real space cut-off of 1.0 nm and SHAKE algorithm was used to constrain all bonds involving hydrogen atoms. The structural figures were rendered using Visual Molecular Dynamics (VMD) ^35^ and the F1-ATP synthase - CST interactions were identified using PDBsum ^36^.

### Ligand affinity isolation of CST binding protein

Mouse heart was homogenized in cold, freshly prepared buffer (10 mM Hepes, pH 7.4, 0.1 mM EDTA) with a cocktail of protease inhibitors (Protease inhibitor cocktail set III, Calbiochem at 1:100: PMSF 5 mM, benzamidine hydrochloride 50 mg/mL, TLCK [N-Tosyl-Lys Chloromethyl Ketone] hydrochloride 0.1 mM), and centrifuged at 18,000 rpm for 20 min to pellet crude membranes. The membrane preparation was washed once with the same buffer and then solubilized with 1% v/v Triton X-100 in 25 mM Hepes pH 7.4, 100 mM NaCl, 2 mM MgCl_2_, 1 mM KCl and protease inhibitor cocktail as mentioned above, for 2 hr at 4°C, then centrifuged at 18,000 rpm for 1 hr, and the supernatant was used as soluble membrane fraction. Biotinylated human CST was synthesized by placing a biotin residue at the C-terminus of the peptide and a spacer consisting of four amino acids (SGSG): SSMKLSFRARAYGFRGPGPQL-SGSG-Biotin. A ligand affinity column was prepared by incubating 5 mg of biotin-CST with 1 ml of 50% streptavidin agarose resin (Thermo Scientific) in column buffer (25 mM Hepes, pH 7.4, 100 mM NaCl, 2 mM MgCl_2_, 1 mM KCl, 1% Triton and protease inhibitor cocktail) for 1 hr at 4°C.

The streptavidin resin was then washed extensively with the same buffer and incubated with the solubilized heart membranes overnight at 4°C. The resin was packed into a chromatography column, washed with the column buffer containing 0.1% Triton, and bound proteins were eluted with column buffer adjusted down from pH 7.4 to 5.5. Fractions eluted were concentrated by TCA precipitation and analyzed on 10% SDS-PAGE gels, stained with Coomassie blue G-250 (SimplyBlue Safestain; Invitrogen). The resulting ∼53 kDa protein band was excised, subjected to in-gel digestion with trypsin, and the resulting peptides were separated by reverse-phase liquid chromatography followed by Tandem mass spectrometry.

### In-gel digest of the CST-binding protein and LC-tandem-MS/MS analysis

Gel slices were subjected to trypsin digestion, and were analyzed by liquid chromatography (LC, C-18)-tandem-MS/MS with electrospray ionization using QSTAR-Elite hybrid mass spectrometer (AB/MDS Sciex) interfaced to a nanoscale reversed-phase high-pressure liquid chromatography (Tempo) with a 10 cm-180 micron ID glass capillary packed with 5-µm C-18 Zorbax^TM^ beads (Agilent) as described before ^37^.

### Cellular uptake and localization of CST

Primary cardiomyocytes from 2-day old C57BL/6 mice were isolated using a PierceTM Primary Cardiomyocyte isolation kit, according to manufacturer’s protocol. After beating cardiomyocytes were observed (7 days of isolation), they were treated with 5 µM FITC-CST (customized from Bon Opus Biosciences) for 4 hrs. MitoTracker® Deep Red FM (500 nM) was added to the cells 30 minutes prior to harvesting for staining mitochondria. The cells were then fixed with 4% paraformaldehyde and stained with nuclear stain DAPI. The cells were observed in a Keyence Fluorescent Microscope at 40X magnification.

### Determination of mitochondrial membrane potential

C2C12 cells cultured with DMEM plus 10% FBS were seeded into collagen-coated 96-well tissue culture plates in 50% confluence. After a 2-day culture, medium was replaced with the same type of medium with/without 100 nM CST. After four-hour exposure to CST, a final concentration of 100nM Image-iT TMRM reagent (Invitrogen) was added to each well, and then continuously culturing for 30 minutes. Discard the medium, and wash cells with PBS. A 150ul cell lysis buffer was added to each well, simply shaken, and read in a plate-reader at 548/574 nm.

### ATP generation by CST

Primary cardiomyocytes from 2-day old C57BL/6 mice were isolated using a PierceTM Primary Cardiomyocyte isolation kit, according to the manufacturer’s protocol. After beating cardiomyocytes were observed (after 7 days of isolation), they were treated with 1µM CST and its control Saline for 24 hours. ATP synthesis was measured using Firefly Luciferase ATP Assay kit (#28854 from Cell Signaling Technology).

### Statistics

Statistics were performed using GraphPad Prism (version 10.4.0). Data were analyzed either with a Repeated Measure (RM) 2x2 ANOVA, 2-way ANOVA and subjected to post-hoc tests and pairwise comparisons where appropriate. Additionally, we performed unpaired student’s t-test or Mann-Whitney U test depending on Shapiro-Wilk’s test of normality (with results corrected for Levene’s test for Equality of Variances were appropriate). All data are presented as mean ± SEM and significance was assumed when α <0.05.

## Results

### Boolean analysis of RNA Seq data revealed upregulation of 39 fibroblast/cardiomyocyte genes induced by CST

We undertook an omics approach to find out the loss and gain of function of left ventricular genes that are regulated by CST. For the Boolean analysis, we utilized a comprehensive dataset of 25,955 human samples (GSE119087) profiled using Affymetrix Human U133 Plus 2.0 microarrays (**Fig. 1A**). We also performed Boolean analysis of RNA Seq data on the left ventricle of WT, CST-KO, and CST-KO+CST mice. *SPARC* (Secreted Protein Acidic and Cysteine Rich), a seed gene known to be expressed in fibroblasts ^38^ and myocytes ^39^, was selected for the analysis. Boolean implication analysis [e.g., *SPARC* high => *TYROBP* (Transmembrane Immune Signaling Adaptor *TYROBP*) high] in blood and leukemia samples indicated that *SPARC* might also be expressed in some immune cell types (i.e. that express *TYROBP*). Therefore, we explored Boolean implications where *SPARC* low => X low to identify genes specific to fibroblasts and myocytes.

**Fig. 1.**
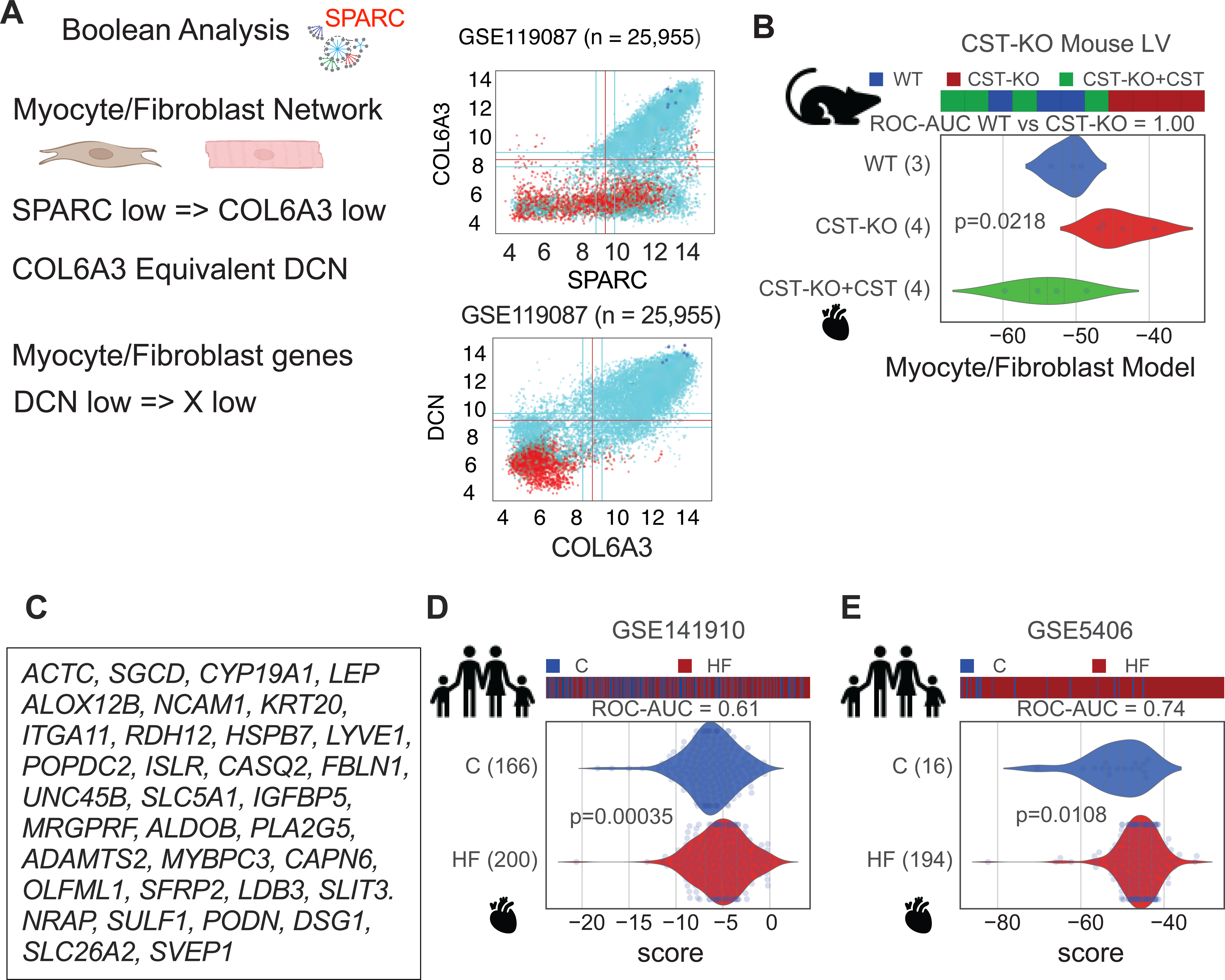
Boolean analysis of RNA Seq data showing CST-induced gene signature focused on cardiomyocyte and fibroblast-specific genes. (A) Boolean Implication analysis to focus on myocyte and fibroblast-specific genes. (B&C) Machine learning using CST-KO RNASeq dataset to identify 39 genes (MyoFibro signature) associated with upregulation in untreated CST-KO mice and down-regulated upon CST treatment. (D&E) Composite score of the MyoFibro significantly distinguished disease states (Control vs heart failure) in human heart samples.

*COL6A3* (Collagen Type VI alpha 3 Chain) was identified as a candidate gene that is not expressed in immune cells but highly expressed in fibroblasts and myocytes. Further analysis of Boolean equivalences involving *COL6A3* led to the identification of *DCN* (Decorin), which showed a Boolean equivalence relationship with *COL6A3* and was highly conserved between humans and mice. This relationship was exemplified by the robust Boolean implication *SPARC* low => *DCN* low. Based on these findings, we hypothesized that genes satisfying the Boolean implication *DCN* low => X low are specifically expressed in fibroblasts and myocytes (**Fig. 1A**).

We filtered genes that were characteristically upregulated in the left ventricles of untreated CST-KO mice and downregulated upon CST treatment. This analysis identified 39 genes constituting the “MyoFibro gene signature,” which are specific to fibroblasts and myocytes (**Fig. 1B&C**): *ACTC* (Actin Alpha Cardiac Muscle 1), *SGCD* (Sarcoglycan Delta), *CYP19A1* (Cytochrome P450 Gamily 19 Subfamily A Member 1), *LEP* (Leptin), *ALOX12B* (Arachidonate 12-Lipoxygenase, 12R Type), *NCAM1* (Neural Cell Adhesion Molecule 1), *KRT20* (Keratin 20), ITGA11 (Integrin Subunit Alpha 11), *RDH12* (Retinol Dehydrogenase 12), *HSPB7* (Heat Shock Protein Family B (Small) Member 7), *LYVE1* (Lymphatic Vessel Endothelial Hyaluronan Receptor 1), *POPDC2* (Popeye Domain Containing 2), *ISLR* (Immunoglobulin Superfamily Containing Leucine Rich Repeat), *CASQ2* (Calsequestrin 2), *FBLN1* (Fibulin 1), *UNC45B* (Unc-45 Myosin Chaperone B), *SLC5A1* (Solute Carrier Family 5 Member 1), *IGFBP* (Insulin Like Growth Factor Binding Protein 5), *MRGPRF* (MAS Related GPR Family Member F), *ALDOB* (Aldolase, Fructose-Bisphosphate B), *PLA2G5* (Phospholipase A2 Group V), *ADAMTS2* (ADAM Metallopeptidase with Thrombospondin Type 1 Motif 2), *MYBPC3* (Myosin Binding Protein C3), *CAPN6* (Calpain 6), *OLFML1* (Olfactomedin Like 1), *SFRP2* (Secreted Frizzled Related Protein 2), *LDB3* (LIM Domain Binding 3), *SLIT3* (Slit Guidance Ligand 3), *NRAP* (Nebulin Related Anchoring Protein), *SULF1* (Sulfatase 1), *PODN* (Podocan), *DSG1* (Desmoglein 1), *SLC26A2* (Solute Carrier Family 26 Member 2), and *SVEP1* (Sushi, von Willebrand Factor Typ1 A, EGF and Pentraxin domain containing 1). Composite score analysis revealed significant upregulation of the MyoFibro gene signature in untreated CST-KO mice (**Fig. 1B**). Furthermore, this signature was significantly upregulated in the heart tissues of heart failure patients (**Fig. 1D&E**).

### Boolean analysis of RNA Seq data identified 34 macrophage genes downregulated in CST-KO mice

Following a similar approach, we also identified macrophage-specific gene signatures (Mac1 and Mac2) using Boolean analysis (**Fig. 2**). Using the same dataset of 25,955 human samples (GSE119087), we performed Boolean implication analysis, which previously identified two universal macrophage genes, *TYROBP* and *FCER1G* (PMID: 32322218). We used the Boolean implication *TYROBP* low => X low to identify additional macrophage-specific genes (**Fig. 2A**). Genes that were downregulated in the left ventricles of untreated CST-KO mice and upregulated upon CST treatment were used to define the so-called “Mac1 gene signature”, comprising 34 genes potentially expressed in various macrophage populations (**Fig. 2B&C**): *IRAK3* (Interleukin 1 Receptor Associated Kinase 3), *RASGRP4* (RAS Guanyl Releasing Protein 4), *CCL18* (C-C Motif Chemokine Ligand 18), *FPR2* (Formyl Peptide Receptor 2), *RDH16* (Retinol Dehydrogenase 16), *SIGLEC10* (Sialic Acid Binding Ig like Lectin 10), *FABP4* (Fatty Acid Binding Protein 4), *ADH1B* (Alcohol Dehydrogenase 1B (Class I), beta Polypeptide), *FPR3* (Formyl Peptide Receptor 3), *MPL* (MPL Proto-Oncogene, Thrombopoietin Receptor), *IFNG* (Interferon Gamma), *CD8A* (CD8 Subunit Alpha), *HDC* (Histidine Decarboxylase), *KLRB1* (Killer Cell Lectin Like Receptor B1), *MS4A14* (Membrane Spanning 4-Domains A14), *ACRBP* (Acrosin binding protein), MS4A3 (Membrane Spanning 4-Domains A3), *CCL7* (C-C-Motif Chemokine Ligand 7), *SRGN* (Serglycin), *IL17RA* (Interleukin 17 Receptor A), *OR2W3* (Olfactory Receptor Family 2 Subfamily W Member 3), *CD300C* (CD300c Molecule), *MS4A4A* (Membrane Spanning 4-Domains A4A), *CLEC4E* (C-Type Lectin Domain Family 4 Member E), *STAB1* (Stabilin 1), *FES* (FES Proto-Oncogene, Tyrosine Kinase), *CCL8* (C-C Motif Chemokine Ligand 8), *CRP* (C-Reactive Protein), *CAMP* (Cathelicidin Antimicrobial Peptide), *IL18RAP* (Interleukin 18 Receptor Accessory Protein), *FGR* (FGR Proto-Oncogene, Src Family Tyrosine Kinase), and *CLEC12B* (C-Type Lectin Domain Family 12 Member B). Composite score analysis revealed significant downregulation of the Mac1 gene signature in untreated CST-KO mice (**Fig. 2B**), with further validation showing significant downregulation in heart tissues of heart failure patients (**Fig. 2D&E**).

**Fig. 2.**
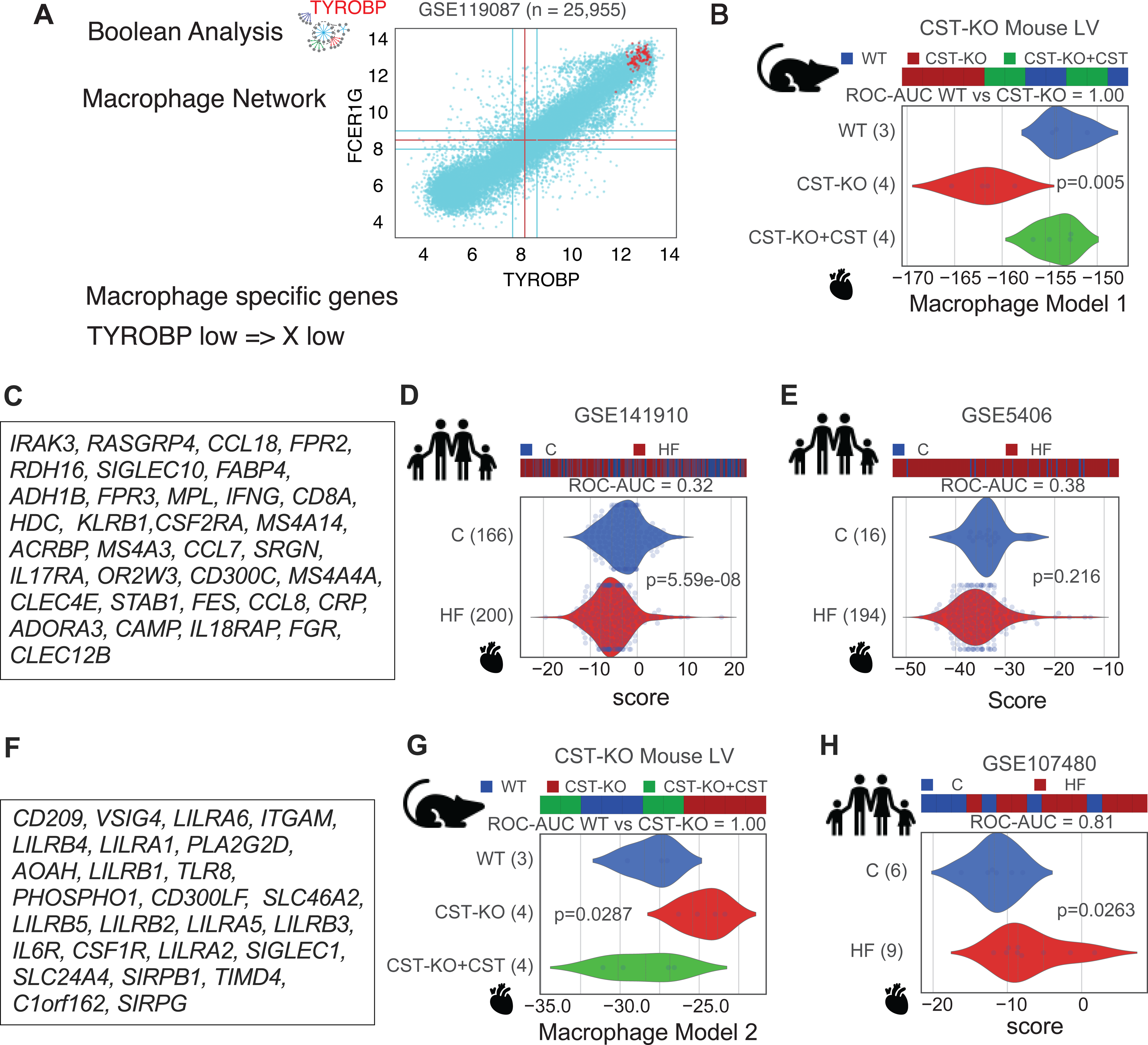
Boolean analysis of RNA Seq data showing CST-induced gene signature focused on macrophage-specific genes. (**A**) Boolean Implication analysis was used to focus on macrophage-specific genes. (**B&C**) Machine learning using CST-KO RNASeq dataset to identify 34 genes (Mac1 signature) associated with downregulation in untreated CST-KO mice and up-regulated upon CST treatment. (**D&E**) Composite score of the Mac1 was able to significantly distinguish disease states (Control vs heart failure) in human heart samples (**F&G**). Machine learning using CST-KO RNASeq dataset to identify 26 genes (Mac2 signature) associated with upregulation in untreated CST-KO mice and downregulated upon CST treatment. (**G**) Composite signature of Mac2 was significantly upregulated in untreated CST-KO mice. (**H**) Composite score of the Mac2 was able to significantly distinguish disease states (Control vs heart failure) in human heart samples.

### Boolean analysis of RNA Seq data recognized upregulation of 26 macrophage genes in CST-KO mice

Additionally, we identified a second macrophage-specific signature, called “Mac2”, comprising 26 genes that were upregulated in untreated CST-KO mice (**Fig. 2F&G**): *CD209* (CD209 molecule), *VSIG4* (V-Set and Immunoglobulin Domain Containing 4), *LILRA6* (Leukocyte Immunoglobulin Like Receptor A6), *ITGAM* (Integrin Subunit Alpha M), *LILRB4* (Leukocyte Immunoglobulin Like Receptor B4), *LILRA1* (Leukocyte Immunoglobulin Like Receptor A1), *PLA2G2D* (Phospholipase A2 Group IID), *LILRB1* (Leukocyte Immunoglobulin Like Receptor B1), *TLR8* (Toll Like Receptor 8), *PHOSPHO1* (Phosphoethanolamine/Phosphocholine Phosphatase 1), *CD300LF* (CD300 Molecule Like Family Member F), *SLC46A2* (Solute Carrier Family 46 Member 2), *LILRB5* (Leukocyte Immunoglobulin Like Receptor B5), *LILRB2* (Leukocyte Immunoglobulin Like Receptor B2), *LILRA5* (Leukocyte Immunoglobulin Like Receptor A5), *LILRB3* (Leukocyte Immunoglobulin Like Receptor B3), *IL6R* (Interleukin 6 Receptor), *CSF1R* (Colony Stimulating Factor 1 Receptor), *LILRA2* (Leukocyte Immunoglobulin Like Receptor A2), *SIGLEC1* (Sialic Acid Binding Ig Like Lectin 1), *SLC24A4* (Solute Carrier Family 24 Member 4), *SIRPB1* (Signal Regulatory Protein Beta 1), *TIMD4* (T Cell Immunoglobulin and Mucin Domain containing 4), *C1orf162* (Chromosome 1 Open Reading Frame 162), and SIRPG (Signal Regulatory Protein Gamma). This Mac2 gene signature was significantly upregulated in the heart tissues of heart failure patients (**Fig. 2H**).

### Decreased insulin signaling in CST-KO heart

Boolean analysis identified genes (e.g., *ITGAM*, *IFNG*) that are involved in developing insulin resistance. In addition, the metabolic flexibility of the heart is compromised with the development of insulin resistance ^14, 15^. Therefore, we checked whether insulin signaling is affected in CST-KO heart. We found that in CST-KO mice, insulin-induced phosphorylation of AKT was reduced by 31% as compared to saline-treated control (**Fig. 3A&B**). Likewise, insulin-induced phosphorylation of GSK-3β was reduced by ∼35% in CST-KO mice compared to saline-treated control (**Fig. 3C&D**). These findings indicate that insulin signaling in CST-KO mice is markedly compromised in the heart.

**Fig. 3.**
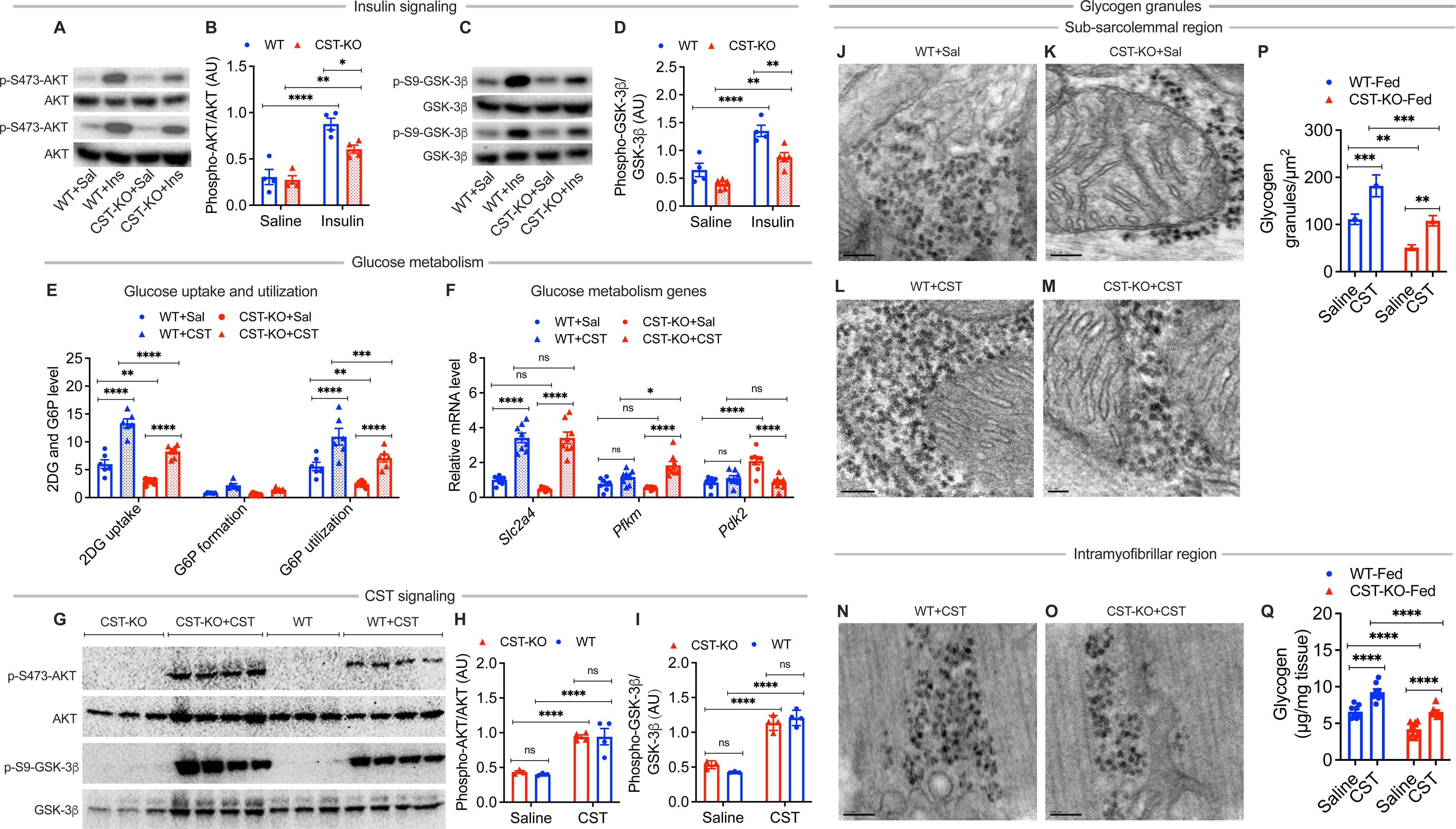
Insulin signaling in wild-type (WT) versus CST-KO heart: (**A**) Western blots showing insulin-induced phosphorylation of AKT. (**B**) Bar graph showing densitometric analysis of the AKT blots shown in (**A**). (**C**) Western blots showing insulin-induced phosphorylation of GSK-3β. (**D**) Bar graph showing densitometric analysis of the GSK-3β blots shown in (**C**). Glucose uptake and utilization in WT and CST-KO heart in response to CST: (**E**) Biochemical determination of 2-deoxy-glucose (2DG) uptake, formation of glucose-6-phosphate (G6P), and utilization of G6P. RT-qPCR data showing expression of genes involved in glucose metabolism in WT and CST-KO heart in response to CST: (**F**) *Slc2a4* gene, solute carrier family 2, member 4 gene (aka *Glut4,* glucose transporter 4); *Pfkm* gene, phosphofructokinase, muscle type; *Pdk2* gene, pyruvate dehydrogenase kinase, isoenzyme 2 gene. CST regulation of proteins involved in insulin signaling: (**G**) Western blots showing CST regulation of phosphorylation of AKT and GSK-3β. (**H**) Densitometric analysis of AKT blots shown in (**G**). (I) Densitometric analysis of GSK-3β blots shown in (**G**). Electron micrographs of the subsarcolemmal region of the heart showing glycogen granules: (**J**) Saline-treated WT heart. (**K**) Saline-treated CST-KO heart. (**L**) CST-treated WT heart. (**M**) CST-treated CST-KO heart. Electron micrographs of the intermyofibrillar region of the heart showing glycogen granules: (**N**) CST-treated WT heart. (**O**) CST-treated CST-KO heart. (**P**) Bar graph showing morphometric analysis of glycogen granules. (**Q**) Bar graph showing biochemical determination of glycogen content in fed WT and CST-KO heart. *p<0.05; **p<0.01; ***p<0.001; ****p<0.0001.

### Decreased glucose metabolism in CST-KO heart

Boolean analysis identified genes (e.g., *ITGAM*, *IFNG, ALDOB, FPR2*) that are involved in glucose metabolism and in the development of insulin resistance. Therefore, we checked how insulin resistance in CST-KO mice affected glucose uptake and utilization in the heart. We found the following: (i) ∼53% decrease in glucose (2DG) uptake in CST-KO mice as compared to WT heart (**Fig. 3E**); (ii) ∼2.2-fold increase in glucose uptake in WT heart in response to CST (**Fig. 3E**); (iii) ∼2.9-fold increase in glucose uptake in CST-KO mice after supplementation with CST (**Fig. 3E**); (iv) no change in G6P formation between and saline or CST-treated WT and CST-KO mice (**Fig. 3E**); (v) ∼57% decrease in G6P utilization in CST-KO heart as compared to WT heart (**Fig. 3E**); (vi) ∼1.95-fold increase in in G6P utilization in WT heart in response to CST (**Fig. 3E**); (vii) ∼2.93-fold increase in G6P utilization in CST-KO mice after supplementation with CST (**Fig. 3E**).

### CST regulation of expression of genes involved glucose metabolism in CST-KO heart

Uptake of glucose into the heart is mediated by Glut1 and Glut4, whereby Glut1 is responsible for most of the basal uptake and Glut4 for most of the stimulus-inducible glucose uptake ^40^. Once inside the cardiomyocyte, glucose undergoes glycolysis, producing 2 pyruvate, 2 ATP, and 2 NADH molecules ^18^. Activation of insulin signaling promotes translocation of Glut4 to increase cardiomyocyte glucose uptake via mechanisms like those described in other cell types ^40, 41^. Therefore, we checked expression of Glut4 gene *Slc2a4* in the presence or absence of the insulin-sensitizing peptide CST. The expression of *Slc2a4* was comparable between WT and CST-KO heart. CST treatment for 2 weeks resulted in ∼3.4-fold and ∼7.1-fold increase in expression of *Slc2a4* gene in WT and CST-KO hearts, respectively (**Fig. 3F**).

In the heart, phosphofructokinase 1 (PFK-1), encoded by *Pfkm* gene, is the rate-limiting enzyme of glycolysis. Like *Slc2a4*, the expression of *Pfkm* gene was also comparable between WT and CST-KO hearts (**Fig. 3F**). Although CST had no effect on the expression of *Pfkm* gene in WT heart, CST caused ∼3.35-fold increase in expression of *Pfkm* gene in CST-KO mice (**Fig. 3F**).

In glucose metabolism, pyruvate dehydrogenase complex (PDC consisting of pyruvate dehydrogenase or E1, dihydrolipoyl acetyltransferase or E2, and dihydrolipoyl dehydrogenase or E3) mediates a major regulatory step, an irreversible reaction of oxidative decarboxylation of pyruvate to acetyl-CoA. PDC activity is tightly regulated using phosphorylation by pyruvate dehydrogenase kinase (PDK1-4) and pyruvate dehydrogenase phosphatases (PDP1 and 2). PDK2 is highly expressed in heart, we checked expression of *Pdk2* gene. Unlike *Slc2a4* and *Pfkm* genes, the expression of the *Pdk2* gene was ∼2.45-fold higher in CST-KO heart compared to WT heart and CST caused ∼58% decrease in expression of *Pdk2* gene in CST-KO heart compared to saline-treated group (**Fig. 3F**).

We also tested for the effects of CST on AKT signaling. Although phosphorylation of AKT at Ser473 was comparable between WT an d CST-KO heart in saline-treated group, CST caused ∼2.19 and ∼2.35-fold phosphorylation of AKT in CST-KO and WT heart respectively (**Fig. 3G**). Like AKT, phosphorylation of GSK-3β at Ser9 was comparable between WT and CST-KO heart in the saline-treated group, CST caused ∼2.13 and ∼2.81-fold phosphorylation of GSK-3β in CST-KO and WT heart respectively (**Fig. 3G**). Unlike insulin (**Fig. 3A&B**), CST was able to overcome insulin resistance and caused comparable phosphorylation of AKT and GSK-3β between CST-KO and WT heart.

### CST increased glycogen content in the heart

Glycogen forms the endogenous cellular energy in the cell and its synthesis is impaired in insulin resistance. Although the heart’s storage capacity for glycogen is limited, its advantage is that it consumes much less oxygen than fatty acids and is readily available for use as fuel. There was a ∼54% decrease in glycogen granules in CST-KO heart compared to WT heart (**Fig. 3J-O, P**). WT and CST-KO heart showed ∼1.64-fold and 2.12-fold increase in glycogen granules respectively in response to CST (**Fig. 3J-O, P**). Fed CST-KO heart, however, showed ∼36.6% less glycogen content in the heart compared to WT heart (**Fig. 3Q**). CST caused ∼1.4-fold increase in glycogen content in fed WT heart as compared to ∼1.56-fold increase in CST-KO heart (**Fig. 3Q**).

### CST inhibited lipid-induced inhibition of insulin-induced glucose uptake in heart

Activation of insulin signaling promotes Glut4 translocation to increase cardiomyocyte glucose uptake. Glut4 translocation is also induced by cardiac muscle contraction. In cardiomyocytes, insulin caused a∼5.9-fold increase in glucose uptake (**Fig. 4A**). Co-incubation with palmitic acid (PA) caused a ∼32% decrease in insulin-simulated glucose uptake (**Fig. 4A**). The presence of CST in the medium not only prevented insulin-stimulated glucose uptake but caused 1.25-fold increase in glucose uptake (**Fig. 4A**). Like PA, co-incubation with ceramide also caused a ∼39% decrease in insulin-simulated glucose uptake (**Fig. 4A**). The presence of CST in the medium not only prevented insulin-stimulated glucose uptake but caused an 1.29-fold increase in glucose uptake (**Fig. 4A**).

**Fig. 4.**
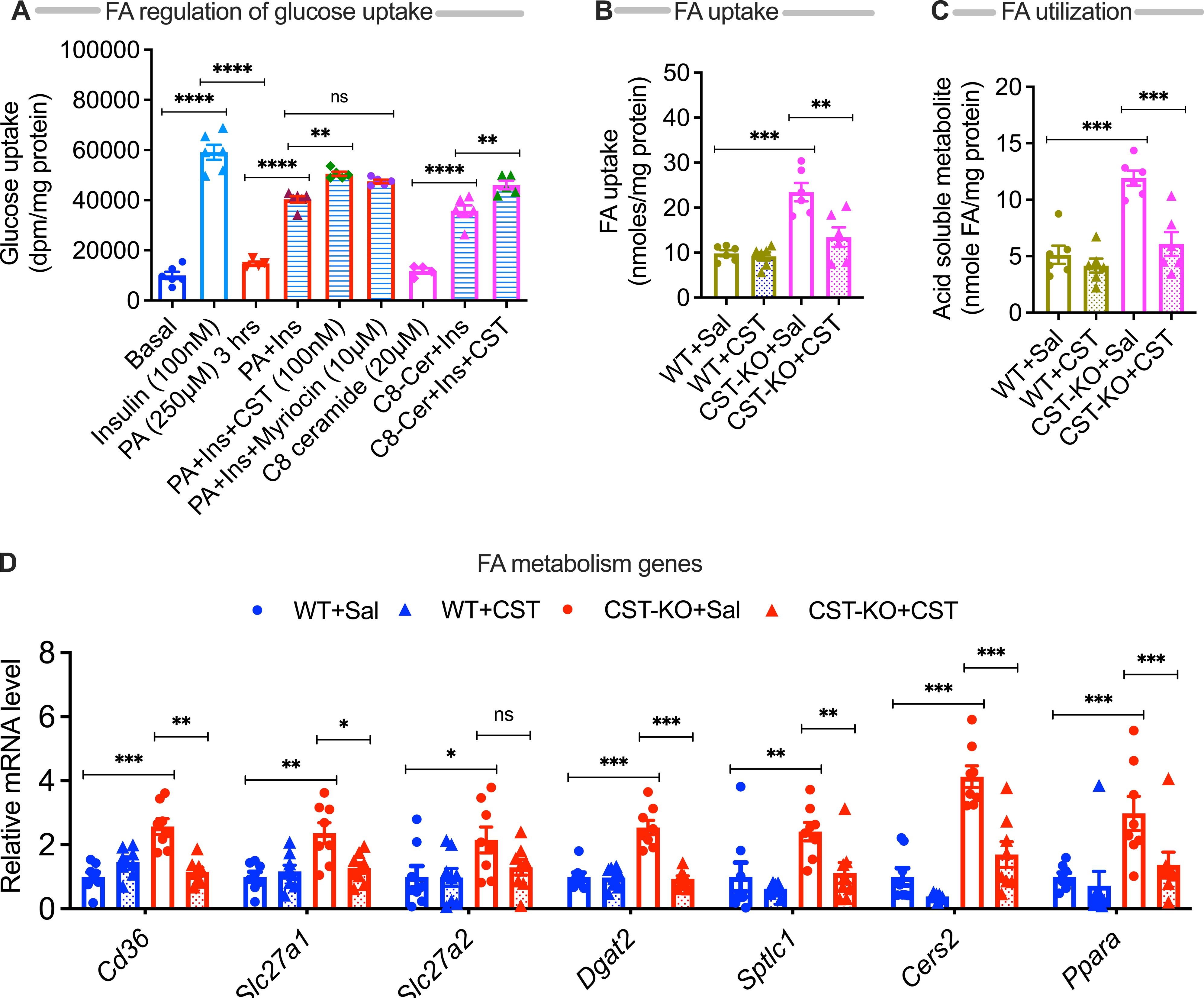
Fatty acid (FA) metabolism in wild-type (WT) versus CST-KO heart. (**A**) Bar graphs showing modulation of insulin-stimulated glucose uptake by FA. (**B**) Bar graphs showing FA update in WT and CST-KO heart in response to CST. (**C**) Bar graphs showing CST regulation of FA utilization in WT and CST-KO heart. (**D**) RT-qPCR data showing expression of genes involved in fat metabolism in WT and CST-KO heart in response to CST: *CD36* gene, Leukocyte differentiation antigen (aka *FAT*, fatty acid translocase); *Slc27a1* gene, solute carrier family 27, member 1 (aka *FATP1*, fatty acid transporter protein 1); *Slc27a2* gene, solute carrier family 27, member 2 (aka *FATP2*, fatty acid transporter protein 2); *Dgat2* gene, diacylglycerol O-acyltransferase 2; *Sptlc1* gene, serine palmitoyltransferase, long-chain base subunit 1; Cers2 gene, ceramide synthase 2; and *Ppara* gene, peroxisome proliferator-activated receptor-alpha. *p<0.05; **p<0.01; ***p<0.001; ****p<0.0001.

### CST decreased expression of FA metabolism genes that were overexpressed in CST-KO mice

Boolean analysis identified genes (e.g., *ITGAM*, *IFNG, ALDOB, FPR2, MS4A4A, PLA2G2D*) that are involved in fat metabolism and in the development of insulin resistance. Although FA can diffuse through cardiomyocyte sarcolemma by a spontaneous diffusional flip-flop process, the bulk of the FA uptake in the myocardium is mediated by the FA transporters CD36 and the plasma membrane fatty acid binding protein (FABP_pm_). FA transport proteins (FATP1 and FATP2 encoded by *Slc27a1* and *Slc27a2*, respectively) also involve FA uptake in the myocardium. FA uptake was ∼2.4-fold higher in CST-KO heart compared to WT heart (**Fig. 4B**). CST treatment caused ∼43% decrease in FA uptake in CST-KO heart (**Fig. 4B**). Consistent with FA uptake (**Fig. 4B**), we found increased expression of *Cd36* (by ∼2.57-fold), *Slc27a1* (∼2.4-fold) and *Slc27a2* (∼2.2-fold) in CST-KO heart (**Fig. 4D**). Supplementation of CST-KO mice with CST decreased expression of *Cd36* (by ∼55%) and *Slc27a1* (by ∼46%) (**Fig. 4D**).

FFAs in healthy tissues are metabolized via beta-oxidation to produce ATP. When calorie consumption exceeds demand, the excess FFAs are packaged into a glycerol backbone to produce inert triglycerides (TGs), which are safely stored in lipid droplets within the cardiomyocytes. The final and the only committed step in the biosynthesis of triglycerides (TGs) is catalyzed by acyl-CoA: diacylglycerol acyltransferase (DGAT) enzymes. Increased expression of *Dgat2* gene (by ∼2.5-fold) in CST-KO heart indicates increased synthesis of TGs in their heart (**Fig. 4D**).

Ceramides, associated with cardiovascular diseases, inhibit nitric oxide synthase, decrease insulin sensitivity, alter mitochondrial bioenergetics, and induce apoptosis and fibrosis. Ceramides are formed by a ubiquitous biosynthetic pathway that starts with the condensation of palmitoyl-CoA and an amino acid (most often serine) to create a sphingoid backbone. Ceramide synthases transfer a FA from acyl-CoA to the sphinganine scaffold, forming dihydroceramide. Ceramide synthase 2 (CERS2), abundantly expressed in the heart, adds very long-chain fatty acids (C20-C26) to the sphinganine scaffold and produces C20-C26 ceramides. Ceramides are essential precursors of most of the complex sphingolipids. Sphingolipid synthesis is catalyzed by the multi-subunit and highly regulated enzyme complex serine palmitoyltransferase (SPT). The function of SPT requires two essential subunits: Serine palmitoyltransferase long chain base subunit 1 (*Sptlc1*) and Serine palmitoyltransferase long chain base subunit 2 (*Sptlc2*). *Sptlc1* catalyzes the initial and rate-limiting step in sphingolipid biosynthesis by condensing L-serine and activated acyl-CoA (most commonly palmitoyl-CoA) to form long-chain bases. CST-KO heart showed increased expression of both *Sptlc1* (by ∼2.4-fold) and Cers2 (by ∼4.1-fold) (**Fig. 4D**). Supplementation of CST-KO mice with CST resulted in marked decrease in expression of both *Sptlc1* (by ∼54%) and *Cers2* (by ∼59%) (**Fig. 4D**).

Peroxisome proliferator-activated receptor-a (PPARα), a transcription factor, plays crucial roles in lipid metabolism in heart. Ubiquitously PPARα-deficient mice showed decreased expression of genes encoding proteins involved in sarcolemmal transport (e.g., CD36 and FATP1), mitochondrial transport (carnitine O-palmitoyltransferase 1, CPT1) and malonyl-CoA decarboxylase, MCD) and mitochondrial (long-chain specific acyl-CoA dehydrogenase, LCAD; medium-chain specific acyl-CoA dehydrogenase, MCAD; and short-chain-specific acyl-CoA dehydrogenase) and peroxisomal (acyl-CoA oxidase) fatty acid oxidation ^42-44^. CST-KO mice displayed ∼3-fold increase in expression of *Ppara* (**Fig. 4D**). Supplementation of CST-KO mice with CST caused ∼54% decrease in expression of *Ppara* (**Fig. 4D**).

### Alteration of sub-cellular structures in heart upon CST knockout

Altered expression of cardiac contractility and mitochondrial health-related genes as identified from RNA sequencing analyses led us to further check any sub-cellular structural changes in myofibrils and mitochondria. Therefore, we sampled left ventricular tissue in diastole after perfusion with cardioplegic solution. We found decreased sarcomere length in CST-KO compared to WT mice in both sub-sarcolemmal and myofibrillar regions (**S-Fig. 1**). We also found evidence of mitophagy in the sub-sarcolemmal mitochondria (SSM) and broken cristae in the inter/intra-myofibrillar mitochondria (IFM) of CST-KO compared to WT mice (**Fig. 5A&B**). CST-KO mice had significantly shorter sarcomeres, and a smaller cristae surface area compared to WT mice (**Fig. 5G&H**). These phenotypes of the CST-KO mice could be reversed by administration of exogenous CST (**Fig. 5C,D,G&H**). In addition, we found lipid droplet (**Fig. 5E**) and mitophagy in the IFM (**Fig. 5F**).

**Fig. 5.**
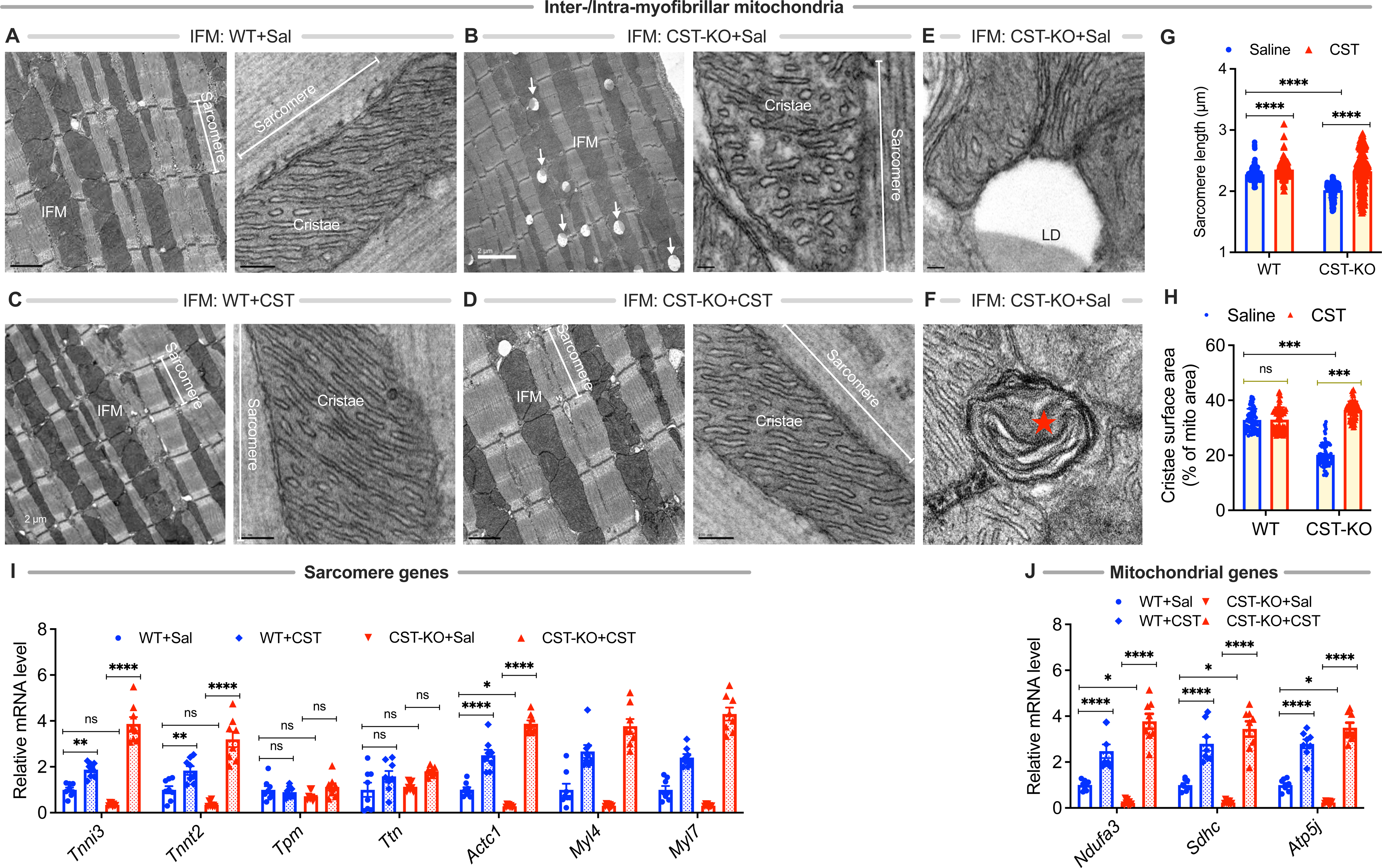
Electron micrographs in the intermyofibrillar regions showing mitochondria in WT and CST-KO heart after treatments with saline or CST. Low and high magnification micrographs in the inter-/intramyofibrillar region: (**A**) saline-treated WT heart showing sarcomere, mitochondria and cristae; (**B**) saline-treated CST-KO heart sarcomere, mitochondria and cristae; (**C**) CST-treated WT heart sarcomere, mitochondria and cristae; (**D**) CST-treated CST-KO heart sarcomere, mitochondria and cristae. (**E**) High magnification IFM showing a lipid droplet. (**F**) High magnification IFM showing mitophagy. (**G**) Morphometric analysis showing sarcomere length. (**H**) Morphometric analysis showing cristae surface area. IFM, inter-/intramyofibrillar mitochondria; LD, lipid droplet; Mp, mitophagy; ZD, Z-disc. ***p<0.001. RT-qPCR data showing expression of sarcomere genes in WT and CST-KO mice after treatment with saline or CST (n=6). (**I**) Note increased expression of the following genes after treatment with CST in WT mice: *Actc1*, actin alpha cardiac muscle; *Myl4,* myosin light chain 4 alkali atrial embryonic and *Myl7,* myosin light chain 7 regulatory and in CST-KO mice: *Tnni3*, troponin I, cardiac; *Tnnt2*, troponin T2, cardiac; *Actc1*. (**J**) Expression of mitochondrial genes in WT and CST-KO mice after treatment with saline or CST (n=6). Note increased expression of the following genes in WT and CST-KO mice: *Ndufa3*, NADH-ubiquinone oxidoreductase subunit A3; *Sdhc,* succinate dehydrogenase complex, flavoprotein subunit A and *Atp5pf*. ATP synthase peripheral stalk, subunit F6 (aka *Atp5j*, ATP synthase, H^+^ transporting mitochondrial FO complex, subunit F6).

### CST regulation of sarcomeric and mitochondrial genes

qPCR studies revealed decreased expression of the following genes: *Tnni3* (by ∼66%), *Tnnt2* (by ∼65%), *Actc1* (by ∼72%), *Myl4/Alc1*, *Myl7/Alc2* (by ∼70%), and *Myh7/Myhcb* (by ∼70%) in CST-KO hearts (**Fig. 5I**). Supplementation of CST-KO mice with CST activated expression of the sarcomeric genes that we have tested here: *Tnni3* (by ∼8-fold), *Tnnt2* (by ∼7.6-fold), *Tpm* (by ∼3.8-fold), *Ttn* (by ∼2.9-fold), *Actc1* (by ∼13-fold), *Myl4* (by ∼12.5-fold), and *Myl7* (by ∼13-fold) (**Fig. 5I**). The expression of the *Tpm1* and *Ttn* genes, however, were comparable between WT and CST-KO hearts (**Fig. 5I**). CST-KO mice showed decreased expression of the following mitochondrial genes: *Ndufa3* (by ∼77%), *Sdhc* (by ∼76%) *Atp5j* (by ∼76%) (**Fig. 5J**). Supplementation of CST-KO mice with CST resulted in increased expression of the mitochondrial genes: *Ndufa3* (by ∼15-fold), *Sdhc* (by ∼13-fold) *Atp5j* (by ∼13-fold) (**Fig. 5J**).

### The 3D-structures of CST and F1-ATP synthase

The time-averaged structure of CST from final 25 ns MD simulation is shown in **Fig. 6A**. The structure is comprised of a metastable antiparallel β-sheet and a random coil (**Fig. 6A**). It was noted that the β-strand persisted for about 60% of the simulation time and converted to random coil or 3_10_ helix at times during the simulation. In the β-sheet conformation, the N-terminal β-strand of CST was formed by the interactions of residues Lys355, Leu356, and Ser357, whereas the C-terminal β-strand was formed by Gly369, Pro370, and Gln371. The modeled structure of mouse F1-ATP synthase was very similar to the crystal structure of bovine F1-ATP synthase that shows an alternate α-β arrangement of three β-subunits and three α-subunits (**Fig. 6B**). Moreover, at the central cavity of this arrangement, there existed an axle made up of γ, δ, and ε-subunits. While the β-subunits are responsible for ATP synthesis, the α-subunits play a critical role in maintaining the structural integrity of F1-ATP synthase.

**Fig. 6.**
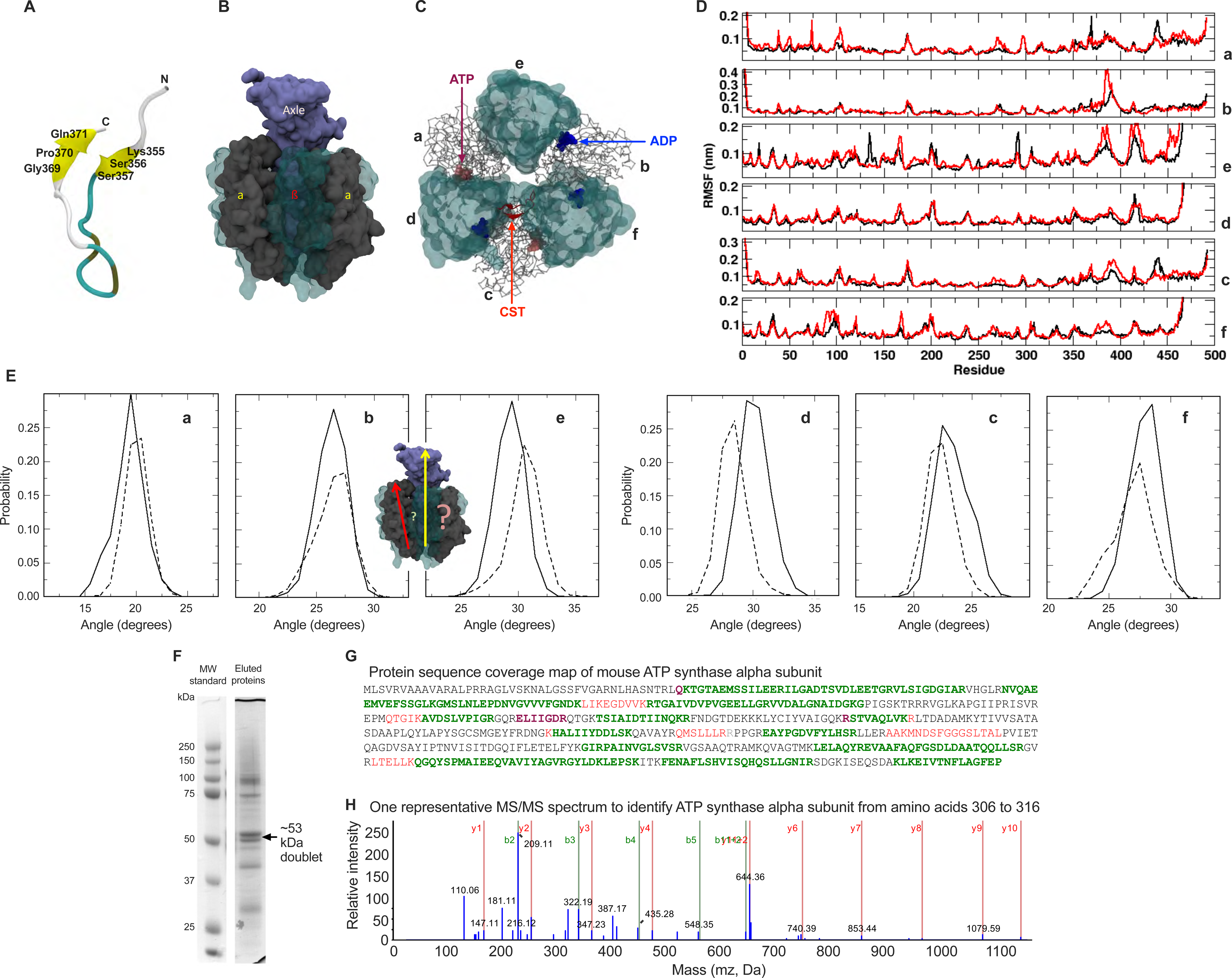
(**A**) Time-averaged structure of CST in cartoon representation. The N and C terminals and some of the important residues are labeled; (**B**) Structure of F1-ATP synthase. The α-subunits are shown in grey, and β-subunits are shown in cyan. The axle is shown in blue. (**C**) Molecular docking of CST to F1-ATP synthase.The a, b, c chains are shown in grey (α-subunits), d, e, f chains are shown in cyan (β-subunits), ATP molecules are shown in magenta, and ADP molecules are shown in blue. Position of CST, which binds to the α-β interface of F1-ATP synthase most effectively (i.e. lowest energy complex) is shown in red. The axle of F1-ATP synthase is hidden for clarity. (**D**) Comparison of root mean squared fluctuations (RMSF) of the residues of F1-ATP synthase subunits. Color scheme – red from simulation data of free F1-ATP synthase (control), black from simulation data of CST-bound F1-ATP synthase at the D-C-F interface. (**E**) Probability distribution of the angle of F1-ATP synthase subunits from the central axle. Results from free and CST-bound F1-ATP synthase simulation are shown by solid and dotted lines, respectively. Identification of a CST binding protein from mouse heart employing affinity chromatography followed by LC-tandem-MS/MS mass spectrometry. (**F**) SDS-PAGE analysis of eluted proteins from biotinylated CST-streptavidin agarose beads. The mouse heart homogenate was subjected to ligand affinity chromatography using biotinylated human catestatin (hCgA_352-372_) peptide as bait. Bound proteins were eluted by lowering pH to 5.5 and analyzed. (**G**) Protein sequence coverage map of mouse ATP synthase subunit α. (**H**) Mouse ATP synthase α-subunit protein sequence coverage map by fragments of a CST-binding protein identified by trypsin-LC-MS/MS; colors are based on peptide confidence: green, peptide has been identified with at least 95% confidence; purple, peptide identified with at least 50% confidence. Regions identified with lower confidence are shown in red. MS/MS spectrum of precursor mass 1286.7 generated after trypsin digestion of CST bound protein.

### CST binding to F1-ATP synthase

The CST-binding data from LC-MS/MS and Time-of-Flight (TOF) has indicated that CST can bind to one of the α-β interfaces of F1-ATP synthase (Table 1). Interestingly, our unbiased protein-protein docking has also shown that CST binds to the α-β interface, specifically at the interface of subunits d, c, and f of F1-ATP synthase (**Fig. 6C**). This is a strong validation of our modeling protocol. A detailed structural analysis revealed that the bound CST is involved in hydrogen bond interactions with Glu341, Tyr381 of D-subunit, and Ser382, Asp315 of F-subunit of the enzyme. Moreover, a large number of hydrophobic interactions of CST residues with the residues of F1-ATP synthase’s c subunit and also with that of D and F subunits were found (Fig. S3). Interestingly, the CST residues Ser352, Arg361, Arg366, and Pro370 were involved in interactions with two neighboring F1-ATP synthase subunits at a time. Such a strong binding of CST can stabilize the α-β interface of F1-ATP synthase, which is necessary for charging ADP to ATP by phosphorylation. To examine the effect of this stabilization on the conformational changes of the opposite interface of subunits a, e, and b of F1-ATP synthase, we performed MD simulations on this docked complex and the results are presented below.

### CST binding at d-c-f interface induces conformational changes in a-e-b interface

MD simulation is being regarded as an important tool in structural biology, as it can explain complex biological phenomena by exploring the conformational changes in the systems at atomic level. Hence, we performed MD simulations on the CST bound F1-ATP synthase complex and also on free F1-ATP synthase as a control. Residue-level fluctuation is commonly used as a metric to decipher the extent of conformational changes in a particular region of a protein. Hence, we measured the Root Mean Squared Fluctuations (RMSF) of F1-ATP synthase residues in both free and CST-bound protein and the results are shown in **Fig. 6D**. As the figure demonstrates, d,c,f subunits of the protein fluctuate less when bound to CST compared to the free state. This again implies that CST binding introduce extra stability to this interface of F1-ATP synthase. On the other hand, the residue-level fluctuations of a, b, and e subunits are observed to increase significantly in CST-bound complex compared to the control system. This suggests that the stability of d-c-f interface due to CST binding induces flexibility to the opposite a-e-b interface of F1-ATP synthase. At this point, it is to be recalled from literature that the inward and outward movements of the constituent subunits of F1-ATP synthase is necessary for ADP binding and ATP release ^45^. To strengthen our result, we also measured the change in angle of each subunit with respect to the central axle in CST-bound and free enzyme. The angle is measured by defining two vectors 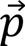 and 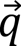, where 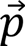 was created by connecting the center of masses (COM) of N-and C-terminal of central axle with the vector directing from N-to C-terminal, while 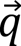 was constructed from the COM of N-terminal β-barrel domain to COM of central β-sheet of each of the protein subunit. Then the angle between the axle and a subunit was calculated by using the following formula:

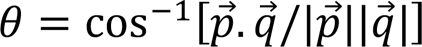

Interestingly, the angle of d, c, f subunits from the central axle was found to decrease, while that of a, b, e subunits increased upon binding of CST to F1-ATP synthase (**Fig. 6E**). This again strengthen the CST-induced increased stability of d-c-f interface and greater flexibility of the opposite a-e-b interface of F1-ATP synthase.

Lastly, we performed “progressive docking” of CST to F1-ATP synthase to examine the possibility of CST binding to other α-β interfaces of F1-ATP synthase. Interestingly, our docking results suggest that four CST peptides can accommodate in the receptor and all the four CST interact at the α-β interfaces of different subunits (**Fig. S4**). Drawing a corollary from our above-mentioned MD simulation results, it is highly probable that the binding of CST at one interface (e.g. a-e-b) will make the opposite interface (d-c-f) flexible. This is in accordance with the fact that each α-β interface of F1-ATP synthase flexes out in succession, due to continuous rotation of the central axle. To identify common interacting residues of CST with different α-β interfaces of F1-ATP synthase, we have also analyzed residue-wise interactions in the docked complexes (**Fig. S5**) and identified seven CST residues, Ser352, Ser353, Arg361, Arg366, Pro370, Gln371, and Leu372, which exhibit common interactions with the α-β interface of F1-ATP synthase. Out of the seven CST residues - Ser352, Arg361, and Arg366 are found to be the common residues at all α-β interfaces, suggesting their significant role in binding.

To conclude, CST has potential to bind to the F1 domain of ATP synthase. Our results from protein-protein docking and MD simulations suggest that CST binding at one interface of F1-ATP synthase can induce significant conformational changes in the opposite interface, as a consequence of which the later flexes away from the central axle. Such conformational changes fit very well with the existing model that proposes the dislocation of a pair of α, β subunits from the central axle for ATP synthesis, while the opposite subunits maintain the stability ^45^. We speculate that in abnormal physiological conditions, such as ischemia, when the rotation of central axle of F1-ATP synthase is completely/partially abolished, CST binding to F1-ATP synthase can maintain the ATP synthesis through a mechanism as proposed above.

### Affinity binding and LC-MS/MS studies confirm CST binding with ATP synthase

We performed experiments to validate ATP synthase binding with CST and to identify other proteins that interact with CST. We used affinity chromatography from murine heart homogenate using biotinylated human CST. CST bound proteins were eluted by lowering the pH from 7.4 to 5.5, followed by analysis with LC-tandem-MS/MS mass spectrometry. The protein bands detected after Coomassie blue staining had M_r_ ∼100, 53, 43, 30 kDa (**Fig. 6F**). When subjected to trypsin digestion and LC-MS/MS analysis, the 53 kDa doublet was identified as the ATP-synthase α- and β-subunit (**Fig. 6F-H; S-Fig. 2**), with 106 spectra spanning ∼74% of the ATP-synthase β-subunit amino acid with overall rank order of one. Tryptic peptides from ATP-synthase subunit α were also identified with 79 spectra and 58% coverage with overall rank order of two (Supplement Table 1). This led us to conclude that CST binds to ATP synthase in murine heart.

In addition, we noted the presence of two ER Ca^2+^-binding proteins (sarcalumenin and calsequestrin), along with trace quantities of commonly observed peptides from tubulin and keratin. Other CST-binding partners are shown in **Table 1**. Interaction between CST and ATP synthase is potentially important not only for ATP synthesis, because ATP synthase is mainly located at the tip of cristae and dimerization of ATP synthase creates a curvature necessary for cristae organization ^30, 31^

### CST penetrates sarcolemma and binds with mitochondria

It has been previously shown that CST can enter neutrophil, bacteria, and fungus ^46^. We used FITC-conjugated CST to find out whether CST could enter cardiomyocytes. Primary cardiomyocytes from C57BL6 WT mice were cultured and treated with FITC-CST. It was observed that CST could penetrate the cardiomyocytes as it co-localized on the mitochondria of the cardiomyocytes that were stained with MitoTracker® Deep Red FM (**Fig. 7A-F**). This co-localization is in line with the computational, molecular simulation and affinity binding studies (**Fig. 6B-H**).

**Fig. 7.**
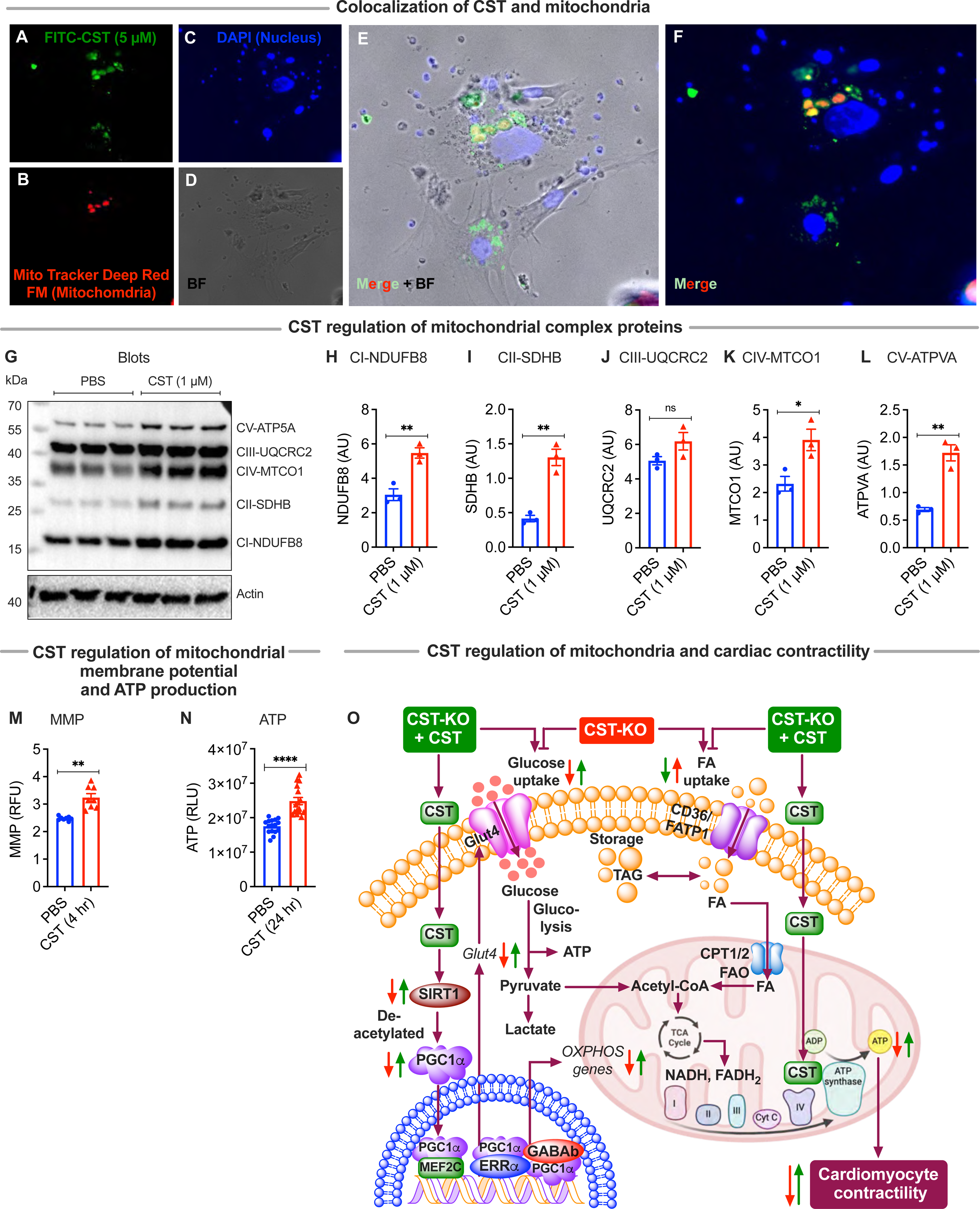
Cell permeable CST co-localizes with mitochondria in cardiomyocytes. (**A**) Incubation of cardiomyocytes with FITC-CST; (**B**) Incubation of cardiomyocytes with Mito Tracker Deep Red FM. (**C**) Incubation of cardiomyocytes with DAPI. (**D**) Bright field photograph. (**E**) Merged plus bright field. (**F**) Merged. Molecular simulation and computer modeling reveal CST binding to α/β subunit of ATP synthase. CST regulation of mitochondrial complex proteins. (**G**) Western blots showing expression of mitochondrial complex proteins. Densitometric analyses of mitochondrial complex proteins. (**H**) Complex I protein: NDUFB8, NADH-Ubiquinone oxidoreductase subunit B8; (**I**) Complex II protein: SDHB, succinate dehydrogenase complex, iron-sulfur subunit B; (**J**) Complex III protein: UQCRC2, Ubiquinol-cytochrome c-Reductase core protein II; (**K**) Complex IV protein: MT-CO1, complex IV, cytochrome c-Oxidase subunit I; and (**L**) Complex V protein: ATP5A, ATP synthase F1 subunit alpha. (**M**) Bar graph showing changes in MMP. (**N**) Bar graph showing changes in ATP concentration. (**O**) Cartoon showing the mechanism of action of CST.

### CST increases mitochondrial function

Our molecular simulation, computational and experimental data showed that CST binds with ATP synthase. The next question we asked was whether such binding modulates mitochondrial function. As a first step in this direction, we looked at the expression of mitochondrial complex proteins in response to CST. We found that with the exception of complex III, CST increased expression of the mitochondrial complex proteins: complex I protein NDUFB8 (by ∼1.79-fold), complex II protein SDHB (by ∼3.17-fold), complex IV protein MTCO1 (by ∼1.68-fold), and complex V protein ATPVA (by ∼2.49-fold) (**Fig. 7G-L**).

It is crucial to maintain a difference in electrical potential between the interior and exterior of the mitochondrial membrane called the mitochondrial membrane potential (MMP; Δψm: generated by proton pumps in Complexes I, II and IV) to sustain the production of ATP. Together with the proton gradient (ΔpH), Δψm forms the transmembrane potential of H^+^, which is harnessed to make ATP. We found that CST caused a ∼31% increase in mitochondrial membrane potential in neonatal cardiomyocytes (**Fig. 7M**), implicating a crucial role that CST plays in optimal functioning of mitochondria.

We then measured ATP content in neonatal cardiomyocytes to find out whether increased Δψm is harnessed to make increased production of ATP. Indeed, we found that CST caused ∼42% increase in production of ATP (**Fig. 7N**). Normally, cells maintain stable levels of intracellular ATP and Δψm, and this stability is believed to be a requisite for normal cell functioning ^47-49^. Thus, CST plays critical roles in maintenance of mitochondrial stability and thereby proper functioning of cardiomyocytes. A schematic diagram showing the mechanism of action of CST is shown in **Fig. 7O**.

## Discussion

CST is considered a “master regulator” of cardiovascular functions as it exerts the following functions: (i) decreases blood pressure in monogenic ^8, 13, 50^ and polygenic ^9, 10, 51^ models of rodent hypertension and in humans ^52^; (ii) decreases both inotropy and lusitropy across different classes of vertebrates: poikilothermic fish ^53^ and amphibia (e.g., frog) ^54^ and homeothermic mammal (e.g., rat) ^55^; (iii) promotes pre-conditioning ^13^ and post-conditioning-induced ^56^ cardioprotection; (iv) maintains baroreflex sensitivity ^57^ and heart rate variability ^58, 59^. In the present study, we found that CST restores metabolic flexibility in diabetic hearts, binds with ATP synthase, and increases MMP and production of ATP.

Conventionally, the heart utilizes both fatty acid and glucose as substrates, with its performance optimal when it can utilize both substrates efficiently ^60^. Substrate utilization changes in response to substrate availability or altered regulation of metabolic pathways ^61-63^. For example, fuel preference switches from glucose and lactate in the fetal heart to predominantly lipids in the adult heart ^62^, while models of cardiac hypertrophy recapitulate the “fetal metabolic profile” with an increased preference for carbohydrate sources ^64, 65^. As CST-KO shows a diabetic heart phenotype, the metabolism shifts away from utilizing glucose, relying almost completely on fatty acids as the energy source. Oxidation of fatty acids has been reported to be metabolically inefficient as it consumes more oxygen molecules than glucose to generate each molecule of ATP ^66^. Thus, it is expected that the diabetic CST-KO heart may suffer from metabolic inflexibility due to reliance on FAs. This metabolic inflexibility partly explain in part the lack of preconditioning-induced cardioprotection in CST-KO mice as reported previously ^13^.

Upon insulin stimulation, CST-KO mice showed a decrease in pAKT and pGSK-3β signals compared to WT mice consistent with lower glucose metabolism in CST-KO hearts. In general, an excess of incomplete fatty acid oxidation contributes to skeletal muscle and cardiac insulin-resistance ^67^. We speculate that the consequence of up-regulated fatty acid uptake and metabolism includes suppression of glucose uptake and utilization in CST-KO hearts through inhibition of pyruvate dehydrogenase (PDH) complex and phosphofructokinase-1 (PFK-1) reactions by excess NADH, acetyl-CoA and citrate respectively ^68, 69^.

Insulin-stimulated AKT and GSK-3β signaling in CST KO hearts suggest a defective Glut4 transporter-mediated glucose uptake in these hearts. Excess acetyl-CoA, derived from increased fatty acid oxidation, inhibits pyruvate oxidation, affecting conversion of excess pyruvate to lactic acid (and associated acidosis). ATP hydrolysis for each molecule of glucose metabolized to lactate (and not oxidized) produces two H^+^ ions (protons) causing acidification ^70^. Acidosis is implicated in contractile dysfunction of the heart ^71^. The excess production of H^+^ may contribute to dysfunction in the failing heart in CST-KO mice. In addition, the increased expressions of genes for the enzymes of ceramide synthesis suggest that ceramide-induced damage could also decrease function in CST-KO hearts.

Gene expression studies show that lack of expression of CST resulted in decreased expression of genes involved in cardiac contraction, including *Tnni3*, *Tnnt2*, *Actc1*, *Myl4*, *Myl7* and *Myh7*. Dilated cardiomyopathy (DCM) is believed to result from a loss of sarcomeric stability or tensile resistance ^72^. The expression of *Tnni3* ^73^ and *Tnnt2* ^73, 74^ genes has been implicated in the development of DCM. Hypertrophic cardiomyopathy (HCM) is believed to arise as a compensation for insufficient contractile force generation ^75^, where ACTC1 plays a significant role ^76^. A novel variant of *ACTC1* gene has been associated with familial atrial septal defect ^77^, while mutation in the *MYL4* gene has been reported to cause familial atrial fibrillation ^78^. Patients with HCM also show increased expression of *MYL7* gene in whole heart tissue ^79^. In human familial HCM, the reduced force generating capacity of sarcomeres is associated with mutation in the *MYH7* gene_80_.

Consistent with perturbation of genes involved in sarcomeric structure and function, ultrastructural studies reveal decreased sarcomere length and decreased A and I bands in CST-KO mice. While increased sarcomere length reflects increased responsiveness to Ca^2+ 81^ and filling of the ventricle ^82^, decreased length has recently been shown to be associated with cardiac dysfunction in intrauterine growth retarded fetuses that persists postnatally ^83^. The length-dependent activation of myofilaments is believed to be due to the reduction of the thick-to-thin filament separation upon an increase in sarcomere length with consequent increase in cross-bridge formation ^84^. Other changes included autophagosomes detected in SSM and broken cristae, in both SSM and IFM in CST-KO mice. In addition, CST-KO mice showed decreased cristae surface area. Mitophagy, a selective autophagy of mitochondria, plays an important role in quality control as it eliminates damaged mitochondria. The observed ultrastructural abnormalities may also contribute to decreased glucose uptake and metabolism, thus both directly and indirectly impacting cardiac function. Decreased glucose uptake and metabolism were consistent with decreased phosphorylation of AKT and GSK-3β in CST-KO compared to WT mice.

Another layer of regulation involves the immune system. Heart-infiltrating macrophages play a crucial role in cardiac inflammation, repair, and remodelling following injury or disease. Our previous studies revealed marked macrophage infiltration in the hearts of CST-KO mice, accompanied by elevated levels of pro-inflammatory cytokines ^13^. While the gene expression profile of macrophages in CST-KO hearts shows a distinct shift compared to WT mice, the data underscore the complexity of this shift. On one hand, several anti-inflammatory genes exhibit reduced expression, including *Irak3*, a negative regulator of TLR signaling ^85^ and the Th2-associated chemokine *Ccl18* ^86^. Conversely, genes associated with pathogen sampling, such as the C-type lectin CD209 and TLR8, show heightened expression. On the other hand, certain pro-inflammatory markers are expressed at lower levels, including the chemokines *Ccl7*and *Ccl8* ^86^ the IL-17A receptor *Il17a*, C-reactive protein *Crp*, and the inflammatory cytokine *Ifng*, whereas *Vsig4*, a molecule known to inhibit pro-inflammatory macrophages ^87^ is upregulated. Intriguingly, we observed upregulation of nine LILR family genes. Since LILR receptors can both inhibit and stimulate antigen presentation and macrophage activation ^88^ these findings further highlight the complex regulatory effects of CST on macrophages.

Our results reinforce the concept that CST is a key immunomodulator in the cardiovascular system. The absence of CST leads to dysregulated macrophage function, potentially contributing to the observed hypertension, cardiac inflammation, and impaired cardioprotection in CST-KO mice ^13^. However, the precise mechanisms by which CST affects macrophages remain an open question. Given the critical role of mitochondrial function in macrophage biology ^88^ it is plausible that CST directly influences macrophage phenotype via mitochondrial modulation, akin to its effects on cardiomyocytes. Importantly, our previous experiments with clodronate-mediated macrophage depletion and bone marrow transfer indicate that CST’s anti-hypertensive effects are partially mediated by its immunosuppressive influence on macrophages, acting as a form of feedback inhibition ^13^.

We conclude that the complex regulation of cardiac heart and mitochondrial structure and function by CST may underlie a key molecular target for metabolic regulation of cardioprotection in the heart. CST may represent a broad therapeutic target for cardiac dysfunction associated with a multitude of diseases (i.e., diabetes, cardiomyopathy, myocardial ischemia-reperfusion, peripheral organ disease induced cardiomyopathy) leading to compromised cardiac structure and metabolism.

## Acknowledgments

Supported by grants from the Veterans Affairs (VA Merit I01BX003934 and VA RR&D SPiRE I21 RX004398-01A1 to SKM; VA Merit BX001963 and VA RCS BX005229 to HHP; VA Merit BX003671 and VA RCS BX006318 to BPH) and the National Institutes of Health (R01 AI163327-01A1 to GG and SKM; R21 AG080246-01 to SKM; R21 AG078635-01A1 to SKM and GG), and the Department of Defense (CDMRP AL210059 (ALSTIA) to BPH; CDMRP AL230115 (ALSTDA) to BPH).

