## Supplemental Figures for "Catestatin improves heart metabolic flexibility by promoting mitochondrial structure and function"

**Table-1**: Catestatin binding protein summary. Top 10 ranked proteins bound to / eluted from the CST affinity column are displayed, as well as the number of peptides identified in each protein. Peptides from the trypsin-digested SDS-PAGE protein bands were analyzed by nano-LC-nano-ESI MS/MS and then identified with Protein Pilot software. Highlighted in red are ER calcium binding proteins, blue = Mitochondrial protein

| Rank | %Cov | Accession | Name | Peptides (95%) |
|---|---|---|---|---|
| 1 | 74.67 | sp\|P56480\|ATPB_MOUSE | ATP synthase subunit beta, mitochondrial OS=Mus musculus GN=Atp5b PE=1 SV=2 | 106 |
| 2 | 58.41 | sp\|Q03265\|ATPA_MOUSE | ATP synthase subunit alpha, mitochondrial OS=Mus musculus GN=Atp5a1 PE=1 SV=1 | 79 |
| 3 | 31.76 | sp\|Q7TQ48\|SRCA_MOUSE | Sarcalumenin OS=Mus musculus GN=Srl PE=1 SV=1 | 26 |
| 4 | 33.41 | tr\|Q52L87\|Q52L87_MOUSE | Tubulin, alpha 1C OS=Mus musculus GN=Tuba1c PE=2 SV=1 | 9 |
| 5 | 23.61 | tr\|Q56A03\|Q56A03_MOUSE | Calsequestrin OS=Mus musculus GN=Casq2 PE=2 SV=1 | 8 |
| 6 | 33.33 | gi\|136429 | RecName: Full=Trypsin; Flags: Precursor | 11 |
| 7 | 29.21 | sp\|Q7TMM9\|TBB2A_MOUSE | Tubulin beta-2A chain OS=Mus musculus GN=Tubb2a PE=1 SV=1 | 2 |
| 8 | 32.81 | sp\|P68368\|TBA4A_MOUSE | Tubulin alpha-4A chain OS=Mus musculus GN=Tuba4a PE=1 SV=1 | 7 |
| 9 | 7.651 | sp\|Q9Z331\|K2C6B_MOUSE | Keratin, type II cytoskeletal 6B OS=Mus musculus GN=Krt6b PE=1 SV=3 | 1 |
| 10 | 7.287 | sp\|Q60936\|ADCK3_MOUSE | Chaperone activity of bc1 complex-like, mitochondrial OS=Mus musculus GN=Adck3 PE=2 SV=2 | 1 |

**Figure legends: Supplemental figures**

**S-Fig. 1. Electron micrographs in the subsarcolemmal regions showing mitochondria in WT and CST-KO heart after treatments with saline or CST. Low and high magnification micrographs in the subsarcolemmal region:** (**A**) saline-treated WT heart showing sarcomere, mitochondria and cristae; (**B**) saline-treated CST-KO heart sarcomere, mitochondria and cristae; (**C**) CST-treated WT heart sarcomere, mitochondria and cristae; (**D**) CST-treated CST-KO heart sarcomere, mitochondria and cristae. (**E**) High magnification SSM showing a lipid droplet. (**F**) High magnification SSM showing mitophagy. (**G**) Morphometric analysis showing sarcomere length. (**H**) Morphometric analysis showing cristae surface area. SSM, Sub-sarcolemmal mitochondria; LD, lipid droplet; Mp, mitophagy; ZD, Z-disc. ***$p<0.001$.

**S-Fig. 2. Identification of a CST binding protein from mouse heart employing affinity chromatography followed by LC-tandem-MS/MS mass spectrometry.** (**A**) Protein sequence coverage map of mouse ATP synthase subunit β. (**B**) Mouse ATP synthase α-subunit protein sequence coverage map by fragments of a CST-binding protein identified by trypsin-LC-MS/MS; colors are based on peptide confidence: green, peptide has been identified with at least 95% confidence; purple, peptide identified with at least 50% confidence. Regions identified with lower confidence are shown in red. MS/MS spectrum of precursor mass 1286.7 generated after trypsin digestion of CST bound protein.

**S-Fig. 3. Residue–level interactions between CST and F1-ATPase.** Interactions of CST with F1-ATPase is depicted. Interactions of three other CST molecules which also can bind to the α-β interface of F1-ATPase, as obtained from progressive protein-protein docking study, are shown subsequently in S-Fig 5. Interestingly, a set of seven CST residues was found to be common in all four CST-F1-ATPase complexes. These residues are highlighted by yellow stars in S-Fig 1 and S-Fig 3. Potential hydrogen bonds between ATPase and CST residues are represented by blue lines and hydrophobic contacts are represented by orange broken lines. The color code given to each residue is based on its nature: positively charged residues in cyan; negatively charged residues in red; neutral charged residues in green; aliphatic residues in grey; aromatic residues in pink; PRO and GLY in orange; CYS in yellow.

**S-Fig. 4. Progressive docking of CST (red) to mouse F1-ATPase.** The α-subunits of F1-ATPase (chain A, B, C) are shown in grey, and the β-subunits (chain D, E, F) are shown in cyan. The location of ATP (magenta) and ADP (blue) in free F1-ATPase crystal structure is also shown.

**S-Fig. 5. Residue–level interactions between CST and F1-ATPase.** Interactions of a) CST-2, b) CST-3, and c) CST-4 with F1-ATPase are depicted. Color scheme is similar to S-Fig 3.

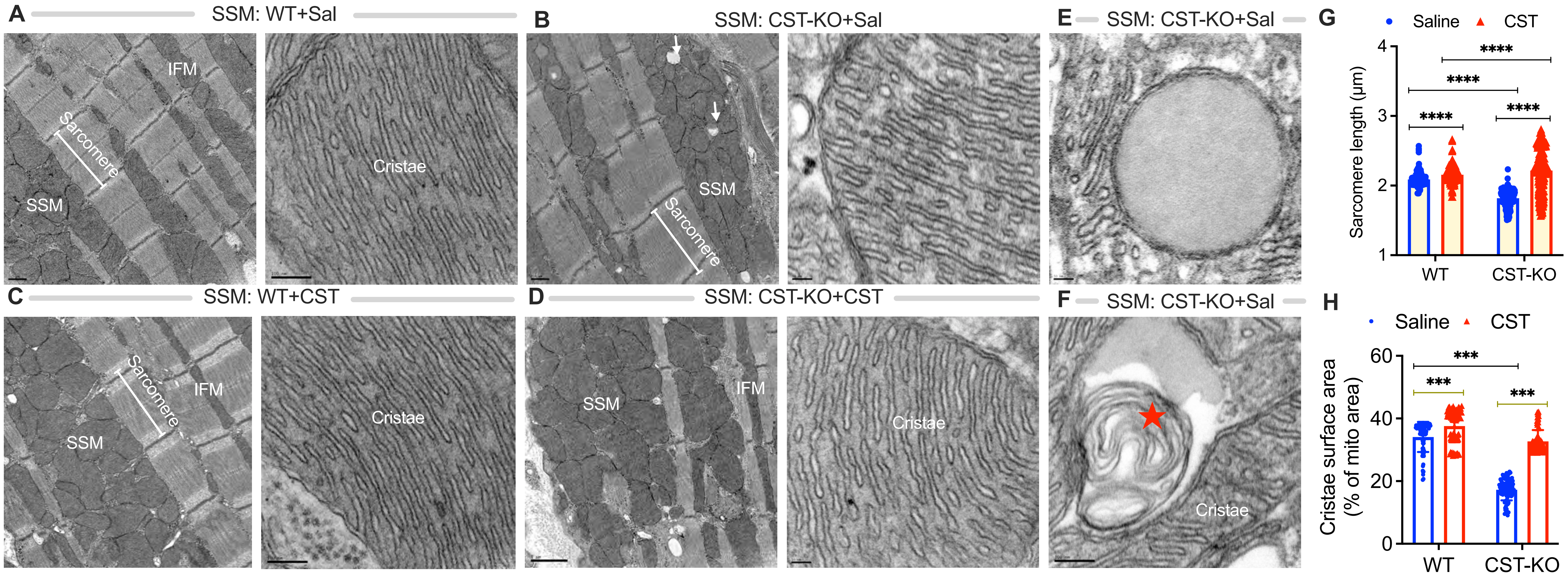

A
SSM: WT+Sal
IFM
Sarcomere
SSM
Cristae
B
SSM: CST-KO+Sal
SSM
Sarcomere
E
SSM: CST-KO+Sal
G
Saline
CST
Sarcomere length (µm)
4
3
2
1
****
****
****
****
WT
CST-KO
C
SSM: WT+CST
Sarcomere
IFM
SSM
Cristae
D
SSM: CST-KO+CST
SSM
IFM
Cristae
F
SSM: CST-KO+Sal
Cristae
H
Saline
CST
Cristae surface area
(% of mito area)
60
40
20
0
***
***
***
WT
CST-KO


S-Fig. 1

**A** Protein sequence coverage map of mouse ATP synthase beta subunit

MLSLVGRVASASASGALRGLSPSAALPQAQLLLRAAPAGVHPARDYAAQASAAPKAGTATGRIVAVIGAVVDVQFDEGLPPILNALEVQGRDS
R**LVLEVAQHLGESTVRTIAMDGTEGLVR**GQK**VLDSGAPIKIPVGPETLGRIMNVIGEPIDER**GPIKTK**QFAPIHAEAPEFIEMSVEQEILVTG**
**IKVVDLLAPYAKGG**K**IGLFGGAGVGKTVLIMELINNVAKAHGGYSVFAGVGER**TR**EGNDLYHEMIESGVINLK**DATSK**VALVYGQMNEPPGAR**
AR**VALTGLTVAEYFRDQEGQDVLLFIDNIFRFTQAGSEVSALLGRIPSAVGYQPTLATDMGTMQER**ITTTK**KGSITSVQAIYVPADDLTDPAP**
**ATTFAHLDATTVLSRAIAELGIYPAVDPLDSTSRIMDPNIVGNEHYDVAR**GVQKILQDYK**SLQDIIAILGMDELSEEDKLTVSR**ARKIQR**FLS**
**QPFQVAEVFTGHMGK**LVPLKETIK**GFQQILAGEYDHLPEQAFYMVGPIEEAVAK**ADKLAEEHGS

**B** One representative MS/MS spectrum to identify ATP synthase beta subunit from amino acids 226 to 239

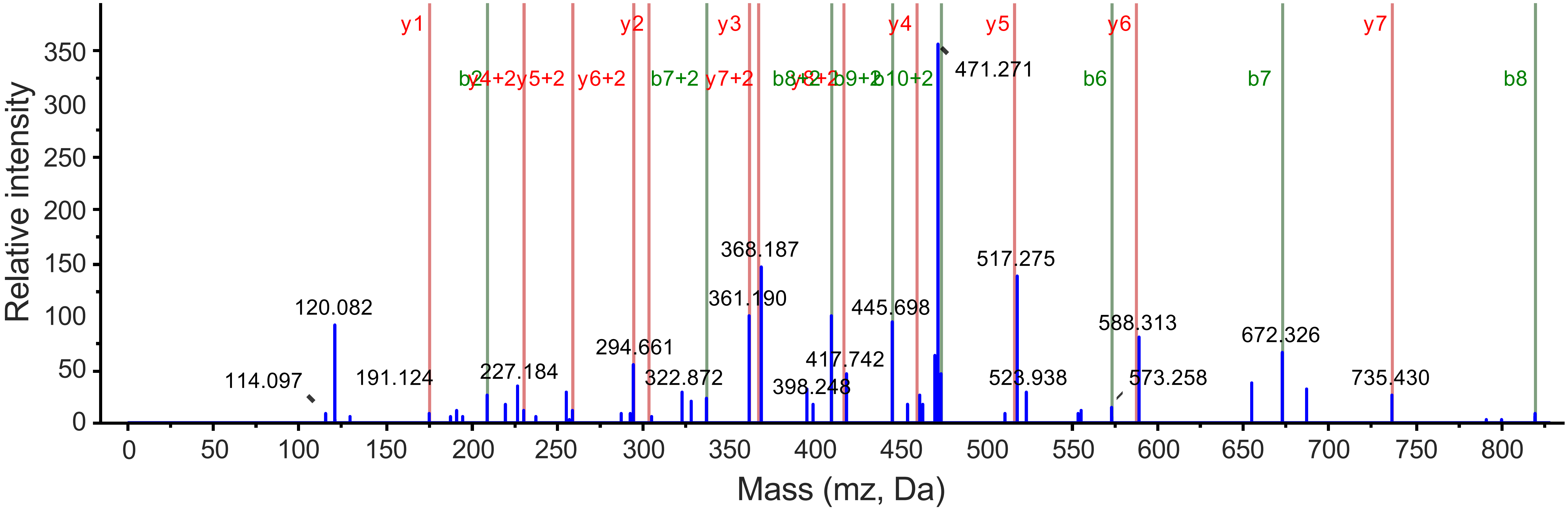


S-Fig. 2

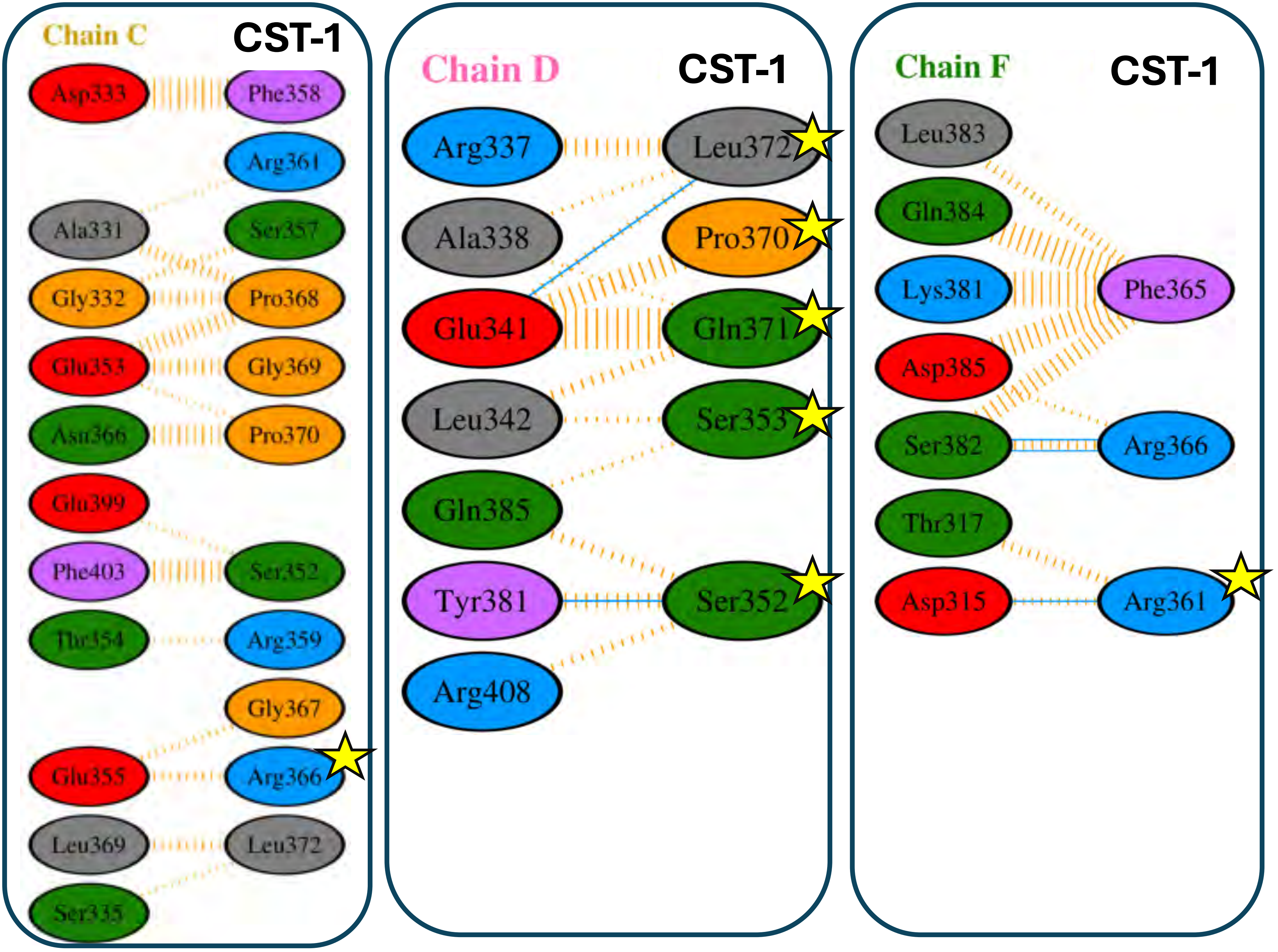

Chain C
CST-1
Asp333
Phe358
Arg361
Ala331
Ser357
Gly332
Pro368
Glu353
Gly369
Asn366
Pro370
Glu399
Phe403
Ser352
Thr354
Arg359
Gly367
Glu355
Arg366
Leu369
Leu372
Ser335
Chain D
CST-1
Arg337
Leu372
Ala338
Pro370
Glu341
Gln371
Leu342
Ser353
Gln385
Tyr381
Ser352
Arg408
Chain F
CST-1
Leu383
Gln384
Lys381
Phe365
Asp385
Ser382
Arg366
Thr317
Asp315
Arg361


S-Fig. 3

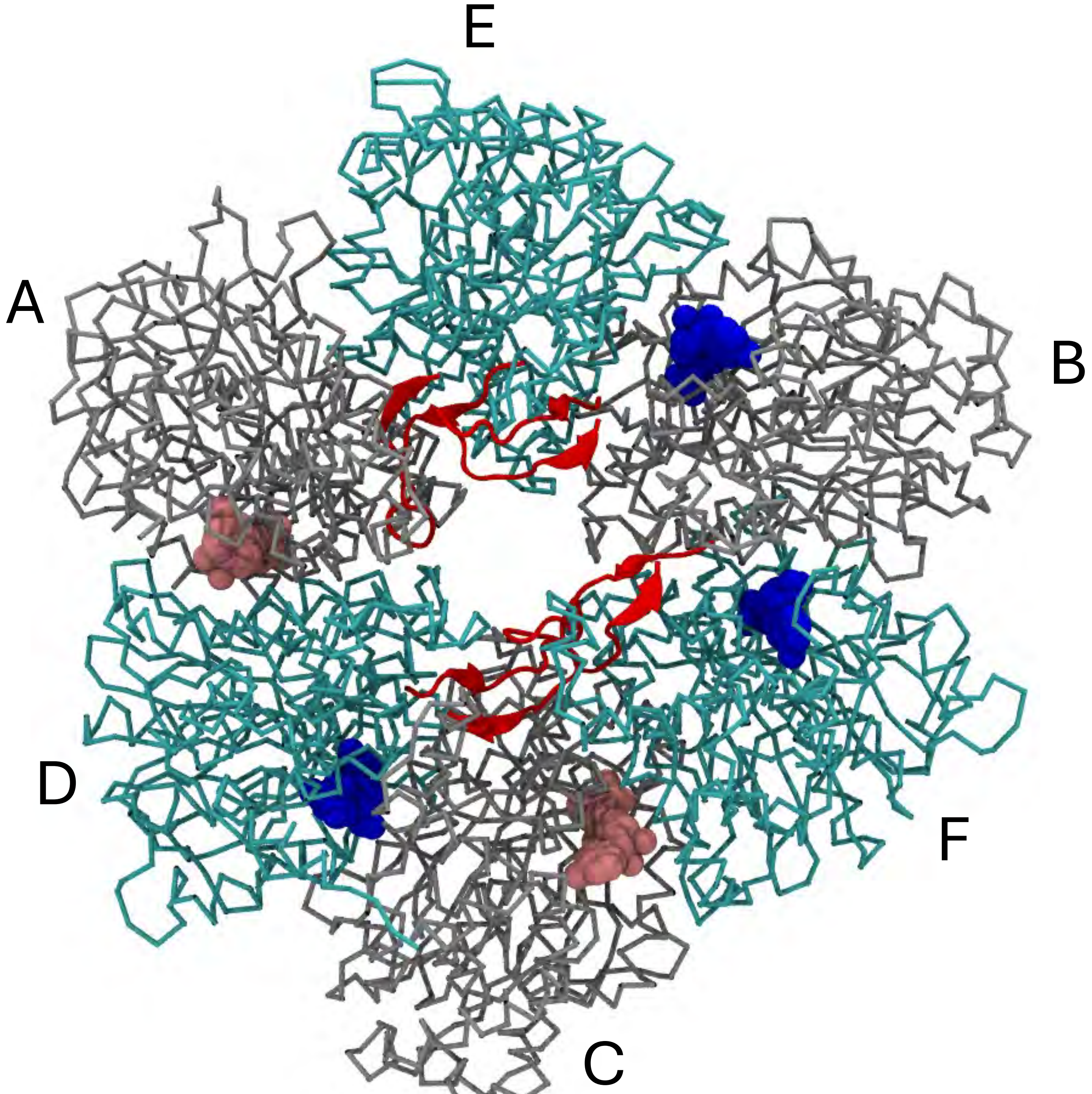


S-Fig. 4

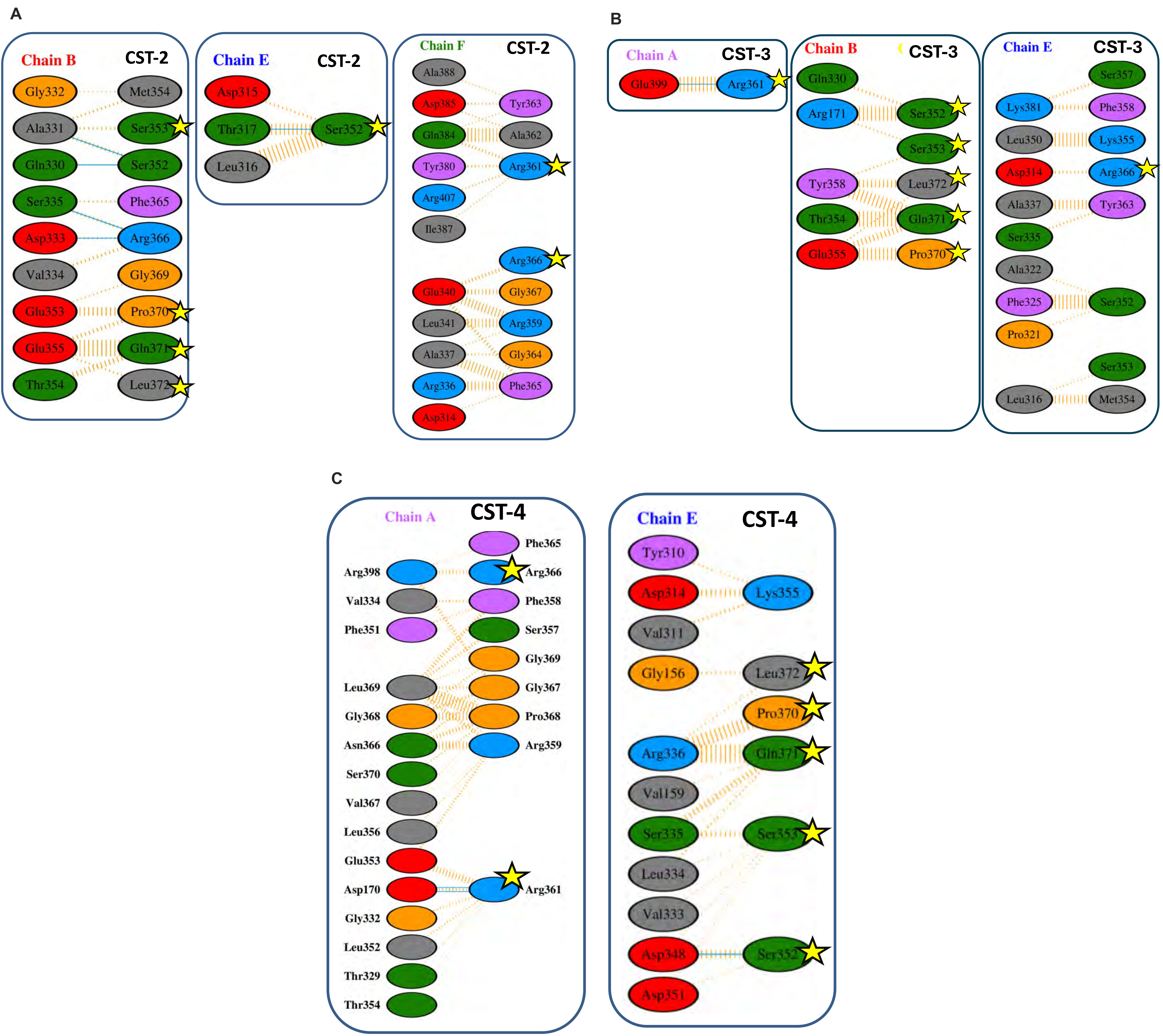
A
Chain B
CST-2
Gly332
Met354
Ala331
Ser353
Gln330
Ser352
Ser335
Phe365
Asp333
Arg366
Val334
Gly369
Glu353
Pro370
Glu355
Gln371
Thr354
Leu372
Chain E
CST-2
Asp315
Thr317
Ser352
Leu316
Chain F
CST-2
Ala388
Asp385
Tyr363
Gln384
Ala362
Tyr380
Arg361
Arg407
Ile387
Arg366
Glu340
Gly367
Leu341
Arg359
Ala337
Gly364
Arg336
Phe365
Asp314
B
Chain A
CST-3
Glu399
Arg361
Chain B
CST-3
Gln330
Arg171
Ser352
Ser353
Tyr358
Leu372
Thr354
Gln371
Glu355
Pro370
Chain E
CST-3
Ser357
Lys381
Phe358
Leu350
Lys355
Asp314
Arg366
Ala337
Tyr363
Ser335
Ala322
Phe325
Ser352
Pro321
Ser353
Leu316
Met354
C
Chain A
CST-4
Phe365
Arg398
Arg366
Val334
Phe358
Phe351
Ser357
Gly369
Leu369
Gly367
Gly368
Pro368
Asn366
Arg359
Ser370
Val367
Leu356
Glu353
Asp170
Arg361
Gly332
Leu352
Thr329
Thr354
Chain E
CST-4
Tyr310
Asp314
Lys355
Val311
Gly156
Leu372
Pro370
Arg336
Gln371
Val159
Ser335
Ser353
Leu334
Val333
Asp348
Ser352
Asp351

S-Fig. 5
